# CellVocal Attributes Transcriptional Dynamics to Spatial Signalling

**DOI:** 10.64898/2026.11.10.765542

**Authors:** Shuai Wang, Jiayin Wang, Xin Lai, Jiayi Wang, Yungang Xu

## Abstract

Understanding how local signalling shapes cell-state transitions requires linking spatial context to transcriptional dynamics. Here we present CellVocal, a framework that integrates spatial RNA velocity with cell-cell communication to decompose cell-state transitions into intrinsic and communication-associated components and further attribute their contributions to signalling modules and neighbouring sender cells. CellVocal recovered complex trajectories and communication perturbations in simulations and reconstructed coherent dynamics and spatially restricted ligand-receptor programmes across the mouse brain, gut and lung. Further analyses linked inferred NCAM1-NCAM1 signalling to lower cell-cycle-associated RNA velocity during embryonic neurogenesis. In melanoma, inferred local MIF-CD74+CD44 signalling corresponded to a predicted decline in the melanocytic programme. Analysis of acute liver failure identified hepatocyte-derived ANGPTL4-SDC1 signalling in association with acetaminophen-related communication enrichment and predicted matrix-remodelling RNA dynamics. By converting static ligand-receptor maps into velocity-constrained networks of signalling influence, CellVocal provides a general strategy for generating spatially resolved hypotheses about how intercellular communication contributes to predicted transcriptional changes.

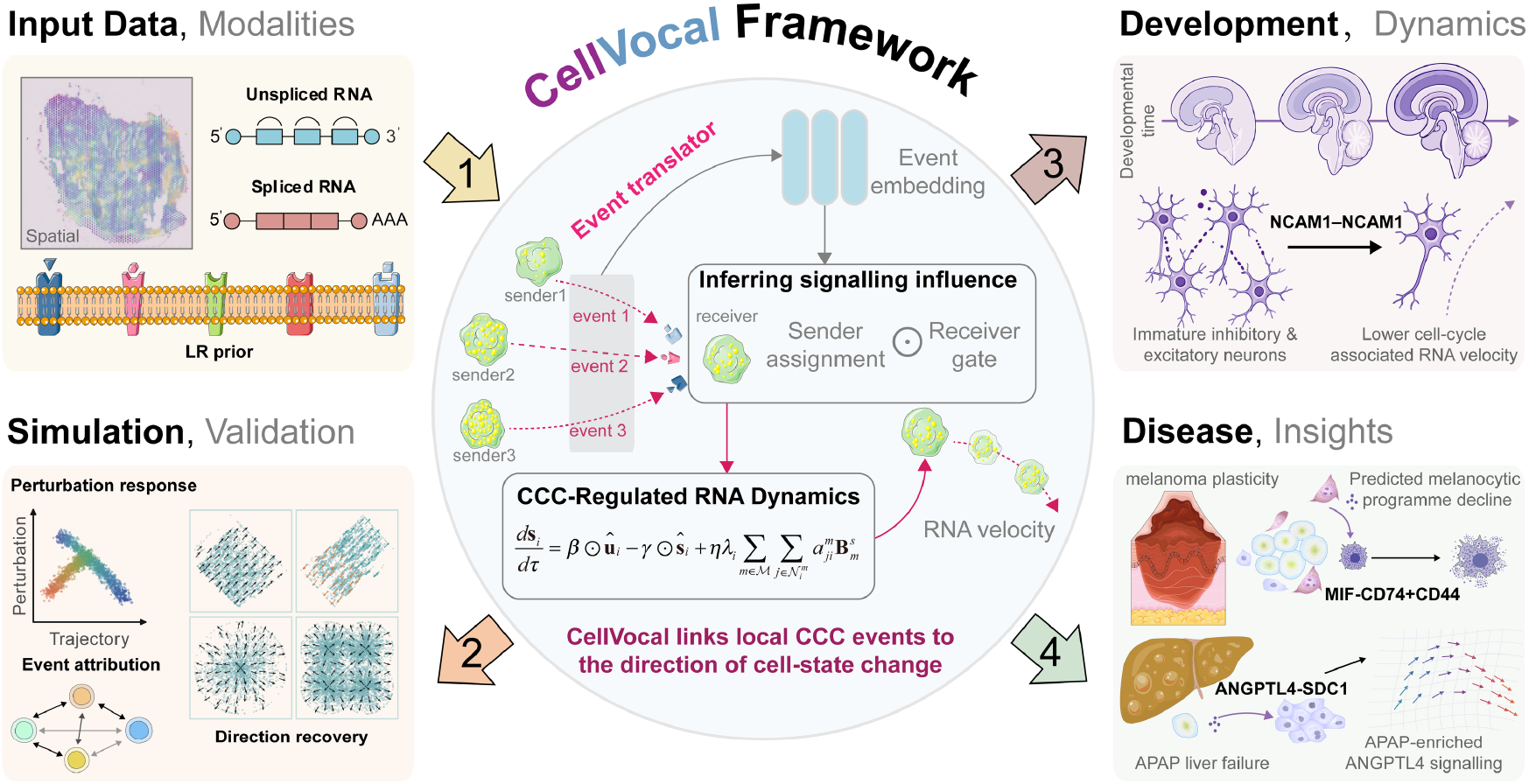

## Introduction

Spatial transcriptomics restores cellular transcriptional states to their native tissue context, enabling tissue architecture, local neighbourhoods, and microenvironmental signals to be resolved within a common spatial coordinate system^1-3^. During dynamic processes such as development, tissue repair and tumour progression, tissue sections are not merely static expression maps but also contain kinetic information describing the continuous cell-state progression across space. RNA velocity estimates the direction and rate of transcriptional change from the relative abundance of unspliced and spliced mRNAs, thereby providing directional information about such cell-state progression in a spatial context^4,5^. Dynamical analysis of spatial transcriptomic data must therefore move beyond characterising the direction of cell-state change towards resolving the origins and regulatory cues of cell-state transitions within the tissue microenvironment.

Over the past several years, RNA velocity methods have increasingly strengthened the characterisation of cell-state progression and transcriptional dynamics. Dynamo extends velocity into a continuous transcriptomic vector field, whereas CellRank combines RNA velocity with transcriptomic similarity to infer directed fate transitions^6,7^. STT and TopoVelo subsequently extended velocity inference to spatial transcriptomic settings, enabling RNA velocity to be integrated with spatial neighbourhoods, tissue topology and local transition structures^8,9^. veloVI, cellDancer and DeepVelo improve velocity inference through uncertainty modelling, cell-specific kinetics and deep-learning-based velocity fields, respectively^10-12^; VeloVAE and Cell2fate further describe complex gene-expression changes using generative models, mixtures of ordinary differential equations and modular transcriptional dynamics^13,14^. VeloCycle, CellRank 2, TIVelo and MultiVelo broaden the applicability of RNA velocity through manifold constraints, multiview fate mapping, trajectory priors and multi-omic coupling^15-18^. These advances enable more robust recovery of cell-state transitions, but most methods still interpret velocity fields primarily as consequences of cell-intrinsic transcriptional dynamics and lack explicit attribution of extracellular signal sources within spatial neighbourhoods. Cell-cell communication (CCC) analysis, by contrast, can identify candidate extracellular signals that may influence cellular states. CellPhoneDB, NicheNet and CellChat infer potential communication networks from ligand-receptor priors, expression patterns and downstream target-gene programmes^19-21^. COMMOT incorporates spatial distance and optimal transport into communication inference, LIANA+ provides a framework for integrating multiple inference methods, and CellNEST further uses spatial graphs and attention mechanisms to resolve local communication structures^22-24^. CellSTIC, which we recently developed, further uses a hierarchical ligand-receptor tree to guide coarse-to-fine graph learning, resolving communication events into traceable multiscale programmes that can be compared across tissues, species and developmental states^25^. However, these approaches generally rank communication events according to co-expression, spatial proximity, or downstream enrichment and thus cannot directly determine whether a candidate signal is aligned with the inferred direction of transcriptional change in the receiver cell.

A critical gap therefore remains between RNA velocity and CCC inference. RNA velocity can recover the direction of cell-state progression but rarely explains which neighbouring sender cells and ligand-receptor modules may drive that progression. Whereas CCC inference can identify candidate communication events but lacks a dynamical constraint derived from RNA velocity. Here we present CellVocal (**V**elocity-**O**riented **C**ommunication **A**ttribution to **L**ocal-senders) for velocity-constrained inference of communication-mediated transcriptional modulation from spatial transcriptomic data. CellVocal centres on RNA velocity and represents changes in receiver-cell states as a coupled process of intrinsic transcriptional dynamics and communication-induced dynamics. For each candidate signalling event, CellVocal estimates signalling influence as the product of the module activation probability in the receiver and the sender’s relative share within that module. CellVocal thereby extends RNA velocity analysis from inference of cell-state transition directions to tracing the communication-associated component of those changes to local signalling modules and sender cells, enabling the systematic investigation of how signals from neighbouring cells participate in cell-state progression within tissues.

## Results

### CellVocal Framework

CellVocal is a communication-aware framework for spatial RNA velocity that resolves how intercellular signals within local microenvironments contribute to cell-state progression. Rather than merely detecting ligand-receptor co-expression between cells, CellVocal decomposes changes in cellular state into intrinsic transcriptional dynamics and communication-induced dynamics, and further traces the communication-induced component to specific signalling modules and neighbouring sender cells. CellVocal thus uses RNA velocity to trace the communication-associated component of receiver-cell transcriptional changes to candidate local signalling events. To establish the dynamical reference frame required for this attribution, the model first integrates unspliced RNA, spliced RNA, spatial coordinates, and optional annotation information. It then uses a spatial neighbourhood graph to encode local RNA states and infer the relative latent time of each spatial location (**Fig. 1a**). This procedure defines both the current position and the direction of progression of each receiver cell within the cell-state landscape, providing the basis for determining whether neighbouring ligand-receptor signals can explain its subsequent cell-state transition.

**Fig. 1.**
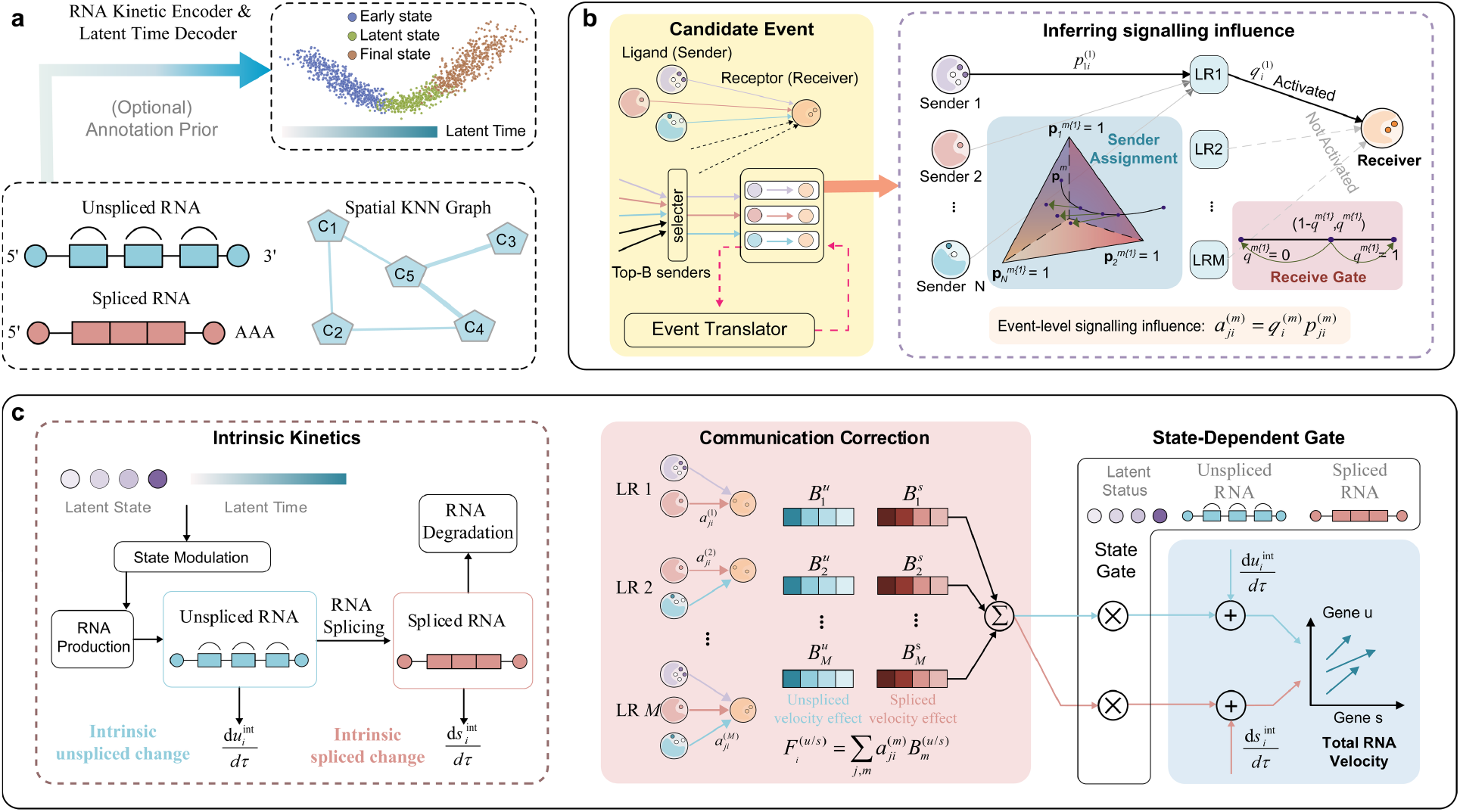
CellVocal framework for communication-aware spatial RNA velocity inference. **a** Unspliced and spliced RNA profiles, optional annotations, and a spatial KNN graph are integrated by the RNA kinetic encoder to infer latent cellular states and relative latent time. **b** Spatially proximal ligand-receptor candidates are selected and translated into communication events; integrating the priors yields an estimate of signalling influence for each sender-receiver event within its module. **c** The RNA velocity decoder combines state-modulated intrinsic RNA dynamics with communication corrections scaled by each event’s inferred influence, gated by the location-specific response strength, to produce the total unspliced and spliced RNA velocity field.

CellVocal subsequently represents communication within the local microenvironment as a process in which multiple senders and signalling modules jointly compete to elicit responses in receiver cells (**Fig. 1b**). Spatial neighbourhoods and ligand-receptor priors are used to define candidate communication events, but signalling influence is inferred from more than ligand-receptor co-expression. To integrate the multiple layers of evidence associated with an individual event, CellVocal uses a cross-attention event translator that encodes each sender-receiver-ligand-receptor event as an event memory containing molecular identity, sender state, receiver state, expression compatibility, and spatial relationships. Event-specific soft queries then adaptively retrieve information from this memory (**Supplementary Fig. 1**). The model can therefore assign context-dependent emphasis to distinct sources of evidence instead of compressing heterogeneous communication features into a single score. CellVocal next estimates receiver-side module activation and within-module sender shares using a two-level conditional categorical flow. At the receiver level, the flow transfers probability mass between inactive and active states to estimate whether a signalling module elicits a response at a given receiving location. At the sender level, the flow redistributes probability mass among the candidate senders associated with the same receiver-module pair, thereby resolving their relative contributions. Because the categorical flows transfer mass over probability simplices under conditional contextual guidance while preserving non-negativity and total mass, the receiver-level distributions distinguish active from inactive modules, while the sender-level distributions capture competition, substitution and cooperation among candidate senders. Analyses of the boundedness and stability of the underlying dynamical system are provided in **Supplementary Note 1**.

Finally, CellVocal uses an RNA velocity decoder to link inferred signalling influence to receiver-cell transcriptional dynamics **(Fig. 1c**). The model first learns state-modulated intrinsic RNA kinetics and subsequently scales module-specific gene effects by each event’s inferred influence to obtain gene-level extrinsic velocity corrections. An event’s inferred influence and its module-specific gene effects jointly determine its contribution to the receiver’s predicted RNA velocity. The outputs of CellVocal therefore comprise not only a spatial communication network but also a set of mechanistically interpretable dynamical readouts, including latent time, total RNA velocity, communication-module activation, and within-module sender shares. By advancing the analysis from “which cells express cognate ligands and receptors” to “which cells, at which locations, contribute to driving the transcriptional-state changes of a receiver”, CellVocal establishes a framework for inferring communication-mediated transcriptional modulation under RNA velocity constraints that resolves the mechanistic relationship between cell-cell communication and cell-state progression in spatially organised tissues. Analyses of the time and space complexity of the complete algorithm, together with its principal computational costs, are provided in **Supplementary Note 2**.

### CellVocal Recovers RNA Velocity-Constrained Cell-Cell Communication in Synthetic and Real Spatial Transcriptomic Data

The central aim of CellVocal is not merely to improve the predictive accuracy of RNA velocity, but to use the inferred direction of transcriptional change as a dynamic reference for tracing communication-associated changes to local signal sources. We therefore constructed three classes of synthetic data spanning direction recovery, communication event attribution, and perturbation responses; detailed generation procedures, parameter settings, and ground-truth definitions are provided in **Supplementary Note 3**. We first tested whether the model could reliably recover latent time and the direction of cell-state progression across different spatial geometries using direction-recovery data comprising linear, bilinear, radial, and multiradial trajectories (**Fig. 2a; Extended Data Fig. 1**). These four trajectory classes represented cell-state progression along a single path, in opposing directions or around multiple spatial centres. We compared CellVocal with six representative spatial or non-spatial RNA-velocity baselines: STT^8^, TopoVelo-GAT and TopoVelo-GCN^9^, scVelo^5^, VeloVAE^13^ and Cell2fate^14^. In the complex simulation containing four radial centres, the latent time inferred by CellVocal closely matched the ground-truth time (Spearman ρ = 0.951; Pearson r = 0.921), with a weighted directional cosine of 0.86 (**Fig. 2b**). Across all four trajectory classes, its mean time correlation, five-step cross-boundary direction correctness (K-CBDir), spatial-velocity consistency and spatial-time consistency were 0.967, 0.941, 0.951 and 0.990, respectively, and it ranked among the top three methods in 11 of the 16 trajectory-metric combinations (**Fig. 2c**). Although some baseline methods achieved higher scores under individual conditions, CellVocal maintained a comparatively stable balance between trajectory direction, local continuity and global temporal ordering, providing the dynamic foundation required to determine whether candidate communication events could explain state changes in receiver cells.

**Fig. 2.**
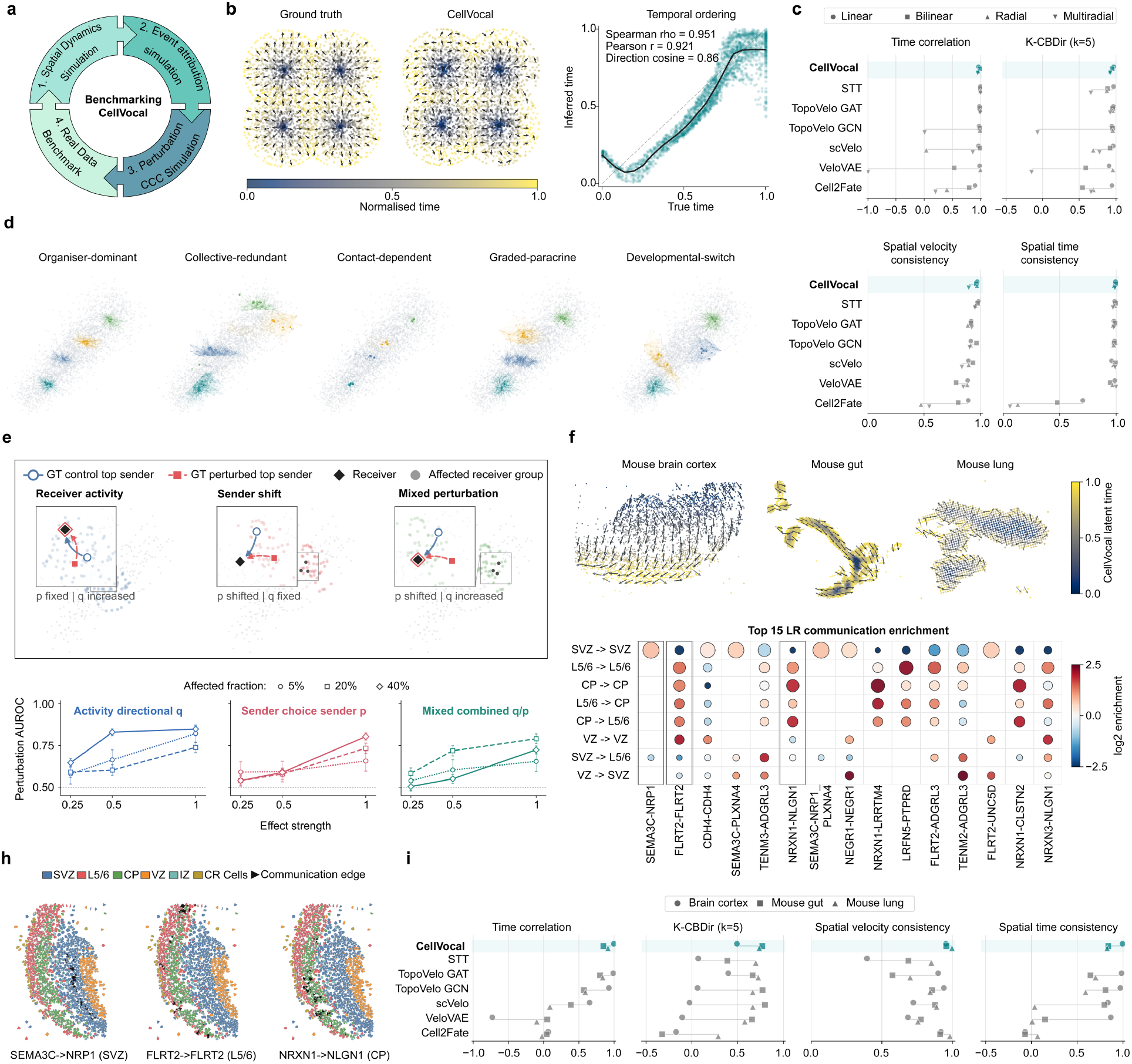
Benchmarking CellVocal in synthetic and real spatial transcriptomic data. **a** Overview of the benchmark design, including direction, CCC, perturbation and real-tissue evaluations. **b** Ground-truth and CellVocal-inferred latent time and velocity field in the multiradial simulation. Arrows indicate spatial velocity; the scatter plot compares inferred and true time, with correlation and direction-cosine scores shown. **c** Comparison of velocity-inference methods across linear, bilinear, radial and multiradial simulations using time correlation, K-CBDir (k = 5), spatial velocity consistency and spatial time consistency. **d** CellVocal-inferred directed CCC networks under five simulated communication mechanisms. Colours denote ligand-receptor pairs and arrows point from sender to receiver. **e** Perturbation AUROC for directional receiver-activity *q*, sender-choice *p*and combined *q*/*p* scores under receiver-activity, sender-shift and mixed perturbations, respectively. **f** CellVocal latent-time and velocity fields in mouse brain cortex, gut and lung. **g** Enrichment of the 15 most active LR pairs across cortical communication routes. Bubble size represents within-LR activity, and colour indicates *log*_2_ enrichment. **h** Spatial communication maps for representative SEMA3C-NRP1, FLRT2-FLRT2 and NRXN1-NLGN1 interactions; cell colours indicate cortical regions. **i** Method comparison in the three mouse tissues using the metrics in c. In c and i, symbols denote datasets or tissues, and horizontal lines show their ranges; higher scores indicate better performance. CCC, cell-cell communication; LR, ligand-receptor.

Having established this directional reference, we next used simulations of event attribution and communication perturbations to test whether CellVocal could recover receiver-side module activation and the relative shares of candidate senders within each module. Five communication mechanisms encompassed distinct patterns of signalling influence, ranging from a single dominant sender to multiple interchangeable senders and from local contact to distance-dependent propagation (**Fig. 2d; Extended Data Fig. 2a, b**). The communication networks recovered by CellVocal retained the principal topological features of each mechanism. Area under the receiver operating characteristic curve (AUROC) values ranged from 0.854 to 0.990 for communication events and from 0.904 to 0.991 for receiver modules, but decreased to 0.545-0.818 when evaluated at the level of exact sender-receiver combinations (**Extended Data Fig. 2c, d**). This distinction is consistent with the hierarchical design of the model: whether a receiver responds to a given signalling module can generally be identified robustly, whereas attributing an exogenous effect to a unique sender is more difficult when several nearby senders carry similar ligand signals. We then used controlled perturbations to determine whether module activation and within-module sender shares could be tracked separately. Activity perturbations changed the strength of the receiver response while preserving sender composition; sender-switching perturbations changed the dominant sender while maintaining overall module activity; and mixed perturbations altered both **(Fig. 2e; Extended Data Fig. 3a**). Across affected-group fractions of 5%, 20% and 40%, the perturbation AUROCs for directional q, sender p and joint q/p scores generally increased with effect size (**Fig. 2e**). The proportions of affected groups, events and differential events exhibited the expected dose structure, with activity perturbations containing only approximately 4.6% differential events (**Extended Data Fig. 3b**). Independent null-model calibration and precision-recall curves showed that event identification was more accurate for sender-switching and mixed perturbations than for activity perturbations, and that the principal scores were generally robust to alternative definitions (**Extended Data Fig. 3c**). The q and p scores for control and perturbed conditions in A09, R09 and M09 further demonstrated the response shifts among differential events (**Extended Data Fig. 3d**). Together, these simulations show that CellVocal can not only establish a reliable reference for cell-state progression, but can also distinguish between two related yet non-equivalent questions: whether the activity of a communication module has changed and whether the source of its signal has shifted.

Finally, we evaluated whether this velocity-constrained communication framework generalised to spatial transcriptomic data from mouse cerebral cortex, intestine and lung (**Fig. 2f**). In comparisons with the same six RNA-velocity baselines, CellVocal recovered velocity fields that varied continuously along the principal tissue structures in all three tissues. It achieved the highest score in 11 of the 12 tissue-metric combinations, with a mean rank of 1.17 (**Fig. 2i; Extended Data Fig. 4**). Within the same dynamic context, communication events scaled by their inferred influence formed distinct ligand-receptor module structures, with 15-nearest-neighbour purities of 0.947-1.000 for event representations in real tissues (**Supplementary Fig. 2**). In the cerebral cortex, SEMA3C-NRP1, FLRT2-FLRT2 and NRXN1-NLGN1 were enriched in the subventricular zone (SVZ), layers 5/6 (L5/6) and cortical plate (CP), respectively (**Fig. 2g, h**).

Ablation experiments further showed that removing the CCC encoder, spatial graph, temporal coupling, or ligand-receptor gating weakened overall performance, with the largest decline observed for the expression-only model (**Supplementary Fig. 3** and **Supplementary Note 5**). By contrast, performance remained broadly stable across the tested ranges of spatial-neighbour number, candidate-event budget, communication-velocity scale and temporal-loss weight (**Supplementary Fig. 4**). Thus, the synthetic and real-tissue results jointly validate the complete CellVocal inference chain: spatial and temporal information first recover the direction of state progression in receiver cells; hierarchical estimation of module activity and sender shares then identifies the signalling modules involved in that change and resolves the neighbouring senders most likely to mediate the exogenous effect within each module. The candidate regulatory signals identified in tissue samples require spatial perturbation or functional experiments to establish their causal roles.

### CellVocal Dissects Spatial Communication to Neuronal RNA Dynamics During Embryonic Neurogenesis

To examine how local adhesion signals relate to neuronal RNA dynamics during development, we analysed the SPTEdU-seq mouse-embryo dataset reported by Niu et al.,^26^ comprising one tissue section each from E12.5, E14.5 and E15.5. Capture sites annotated as immature inhibitory or excitatory neurons were designated as receivers (Fig. 3a). CellVocal estimated incoming signalling influence and gene-response profiles at these sites, enabling candidate neighbourhood signalling events to be examined alongside their predicted transcriptional consequences. The screen assessed the association between incoming signalling influence and the RNA response and compared receivers with high incoming influence against background receivers using *Mki67* and *Top2a* responses, while stage-specific coupling retained the developmental context of each nominated module. Screening 77 ligand-receptor modules prioritised NCAM1-NCAM1 as a candidate regulatory signal. Its incoming influence was consistently associated with a communication-derived cycling-down score across sections; NRXN1-CLSTN2 was the next-ranked candidate (**Fig. 3b**). This model-based prioritisation identified adhesion pathways for subsequent spatial and transcriptional analysis. Local reconstruction traced inferred NCAM1 signalling to individual source-receiver routes in an E14.5 neighbourhood, distinguishing the inferred influence of individual nearby senders sharing the same candidate ligand-receptor pair (**Fig. 3c**). Across neuronal receiver sites in the analysed sections, inferred NCAM1 signalling arose from several source classes and reached both immature neuronal lineages, with broader coverage among excitatory receivers (**Fig. 3d**). The local routes therefore exposed the contributing neighbours beneath the aggregate source-to-lineage network. These adhesion-associated responses have a plausible molecular context: NCAM stimulation can engage PKCα-dependent TRPV2 calcium signalling during neurite outgrowth,^27^ and an NCAM-integrin α7β1 complex transduces Schwann-cell-derived GFRα1 signals during axonal regeneration.^28^ These experiments support the ability of NCAM-containing receptor complexes to translate extracellular cues into neuronal responses, providing a basis for examining how their spatial allocation accompanies RNA dynamics in the developing tissue.

**Fig. 3.**
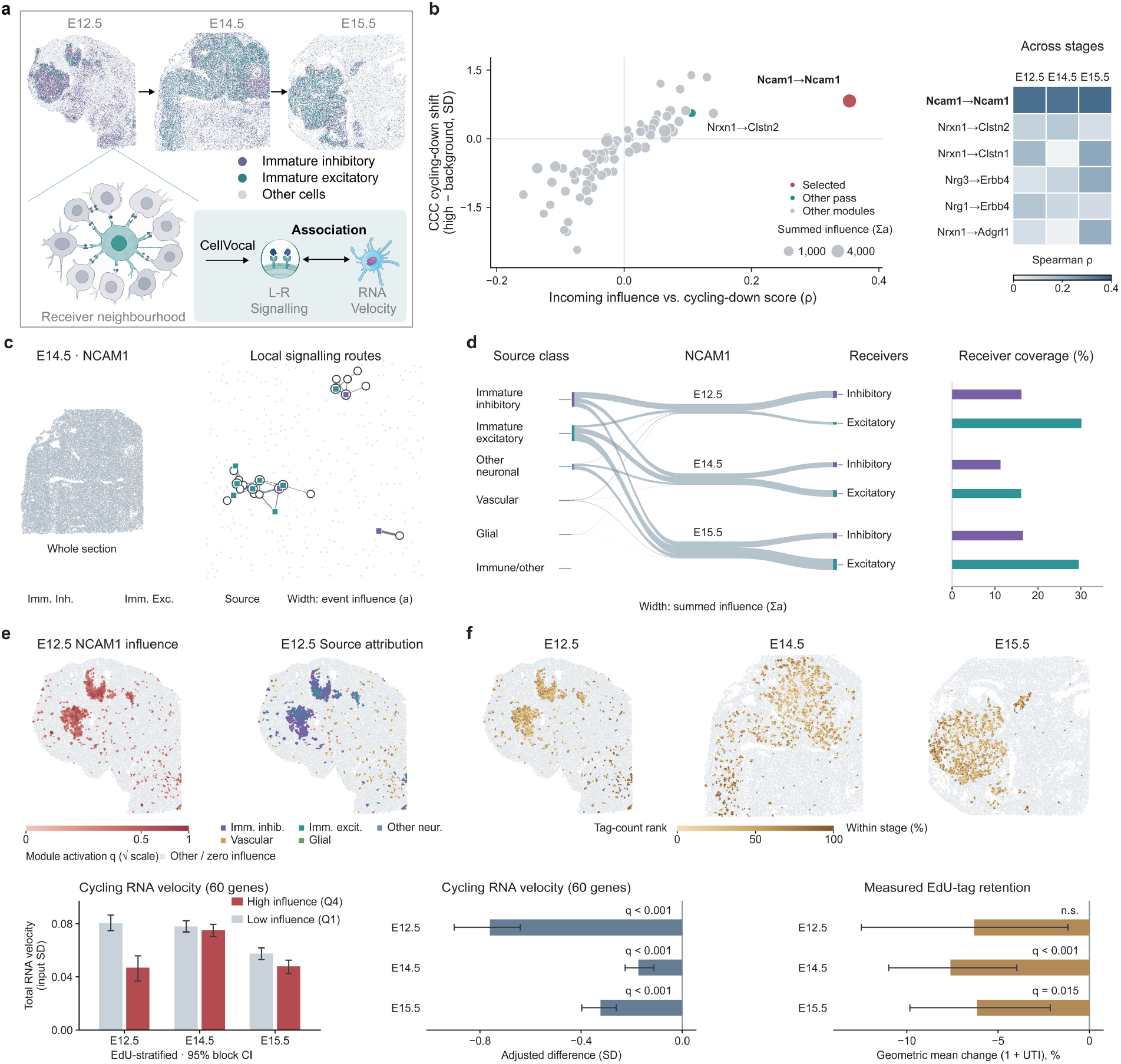
CellVocal resolves NCAM1-associated spatial communication and neuronal RNA dynamics in the mouse embryo. **a** Spatial distributions of immature inhibitory and excitatory neuronal capture sites at E12.5, E14.5 and E15.5. **b** Screening of 77 ligand-receptor modules by the association between incoming signalling influence and the RNA response, together with contrasts between receivers with high incoming influence and background receivers for a communication-associated cycling-down score derived from *Mki67* and *Top2a*. Bubble size indicates summed signalling influence; the heatmap shows stage-specific coupling for the six highest-ranked modules. **c** Local NCAM1 source-receiver routes in E14.5; edge width indicates each event’s inferred signalling influence. **d** NCAM1 signalling influence across source classes, stages and receiver lineages, with the fraction of receivers showing positive incoming NCAM1 influence. **e** Incoming NCAM1 influence and dominant inferred source class in E12.5, and total RNA velocities of a 60-gene cycling programme in lower and upper quartiles of positive incoming NCAM1 influence, averaged across historical EdU-tag/RNA strata. **f** Retained EdU-tag abundance at NCAM1-positive receivers and adjusted cycling-velocity and tag-retention contrasts comparing the median positive estimate of incoming NCAM1 influence with zero, accounting for lineage, RNA depth and spatial blocks. RNA contrasts use outcome standard deviations; EdU contrasts are percentage differences in the geometric mean of one plus tag count. Error bars in e,f show 95% confidence intervals from 2,000 spatial-block bootstrap samples. Tests in f are two-sided with spatial-cluster-robust errors and Benjamini-Hochberg correction across 54 comparisons.

We next examined whether inferred NCAM1 signalling was associated with a change in the receivers’ overall RNA dynamics. The E12.5 maps showed spatial variation in incoming NCAM1 influence and variation in the dominant attributed source class between receiving sites (**Fig. 3e**). This source attribution preserved the connection between each local signalling event and the neuronal population receiving it. Among receivers with positive incoming NCAM1 influence, those in the upper quartile had lower mean total RNA velocity across a 60-gene cycling programme than those in the lower quartile after stratification by historical EdU-tag/RNA ratio. The four EdU strata contributed equally to each group mean. The direction was consistent across sections, with the clearest separation at E12.5 and E15.5 and a smaller contrast at E14.5. Spatial-block bootstrap intervals excluded zero for the high-minus-low difference at E12.5 and E15.5, accounting for local dependence among neighbouring capture sites. Thus, the association extended from the communication-derived screening score to total predicted cycling-associated RNA change. Both groups retained positive programme-average velocities, placing the response in reduced predicted RNA accumulation rather than completed cell-cycle exit. Stratification compared levels of incoming NCAM1 influence within groups of similar retained labelling history, addressing whether the neighbourhood association persisted within that measured aspect of developmental state. Multiple admitted L-R modules could coexist at individual receivers; their gene-specific responses and partly opposing contributions explained how the selected NCAM1-associated signal was integrated with other local signalling contributions (**Extended Data Fig. 5**). The resulting programme readout retained these combined contributions alongside the model’s intrinsic RNA dynamics.

Measured EdU-tag retention provided a complementary view of the populations associated with inferred NCAM1 signalling (**Fig. 3f**). Because EdU was administered at E10.5 and E11.5, the retained tags recorded earlier labelling history. Comparing the median positive estimate of incoming NCAM1 influence with zero, with adjustment for receiver lineage, RNA depth and spatial blocks, identified lower cycling-programme velocity at all three stages. This comparison of positive versus zero incoming NCAM1 influence complemented the within-positive dose contrast by including receivers without inferred NCAM1 signalling. Measured tag retention was also lower in receivers with positive incoming NCAM1 influence at E14.5 and E15.5, whereas the earlier section showed less separation. The paired spatial readouts placed the inferred RNA dynamics in a measured developmental context without treating the tags as direct measurements of subsequent changes in RNA expression. To probe the contribution of the candidate pathways, we removed the NCAM and NRXN communication components with model parameters, measured RNA, and latent states fixed (**Extended Data Fig. 6a,b**). Comparisons used the same receivers with positive incoming influence from the targeted modules within each section, preserving their original spatial positions. Joint removal increased cycling-programme and decreased neuronal-function-programme velocity across all three sections. These broader programme responses accompanied the same respective directions in the two cycling and three neuronal-marker averages. The shifts varied spatially and included increased Mki67 and decreased *Map2* velocity at each stage (**Extended Data Fig. 6c,d**), demonstrating dependence on these communication components within the fitted model. Joint removal also changed velocity direction, with median full-gene angles of 30-32° across sections and corresponding tissue projections (**Extended Data Fig. 6f,g**). Published isogenic NRXN1-deletion organoid contrasts (**GSE228315**) provided complementary expression-level evidence of adhesion-associated neuronal responses, including reduced FOS in several neuronal classes, with developmental and cell-class variation (**Extended Data Fig. 6h**).

### CellVocal Links Local MIF-CD74+CD44 Signalling to Predicted Decline of the Melanocytic Programme in Melanoma

To examine how local signalling relates to changes in tumour-cell identity, we analysed three Visium sections from the mouse melanoma dataset of Karras et al., comprising 7,626 capture spots annotated as malignant.^29^ Melanocytic, neural-crest-like and other malignant states formed interspersed tissue domains, placing spatially distinct neighbourhoods within a heterogeneous tumour cell-state landscape (**Fig. 4a**). Sender and receiver assignments were therefore interpreted at the capture-spot level, consistent with the multicellular resolution of Visium. The inferred latent organisation provided a complementary view of these states within each section (**Extended Data Fig. 7**). We defined predicted melanocytic decline as the negative projection of total RNA velocity onto a fixed 17-gene signature containing *Mitf, Tyr, Dct* and *Pmel*. This readout combined predicted transcript changes into a common direction of lineage-programme change, allowing local communication to be evaluated against a dynamical outcome defined consistently across sections. Among 25 communication modules, MIF-CD74+CD44 combined a positive association between incoming MIF influence and predicted decline with the largest contrast between spots with high incoming influence and background spots (**Fig. 4b**). This prioritisation identified a local signalling candidate associated with the direction of lineage-programme change, rather than simply its current abundance. The joint association remained supported after spatial-rotation testing and correction across candidate modules (P = 0.018), and contrasts between spots with high incoming MIF influence and background spots persisted across a range of thresholds; section-level support was strongest in S4 (**Supplementary Fig. 5a-c**).

CellVocal then resolved the spatial sources and gene-level responses underlying this association. Inferred MIF-CD74+CD44 signalling occupied restricted tissue neighbourhoods, whereas total melanocytic-programme velocity varied across the wider malignant compartment (**Fig. 4c**). At a representative receiver in an S3 neighbourhood, MIF-CD74+CD44, GRN-SORT1 and COL3A1-ADGRG1 signalling contributions converged from several neighbouring senders. CellVocal estimated the relative shares of these senders within each module and decomposed each module’s signed contribution to *Mitf, Tyr, Dct* and *Pmel* RNA dynamics (**Fig. 4d**). The resulting response was gene-specific: MIF-associated contributions differed in sign across the four markers, and other signalling contributions could reinforce or oppose them. Including the remaining modules reconstructed the complete communication-associated response at this receiver. The net response therefore depended on both the strengths of incoming signals and their gene-specific response coefficients, exposing how overlapping signalling modules contributed to different components of the same lineage programme. This decomposition distinguished the presence of a candidate ligand-receptor interaction from its inferred contribution to a particular transcriptional response. Mapping the same routes back into tissue placed the predicted response magnitude attributed to each sender alongside receiver expression and total RNA velocity, showing how multiple local signalling contributions were integrated within a shared spatial niche (**Fig. 4e**). Across the three sections, malignant-labelled sender spots dominated the attributed MIF response magnitude, while a subset of receivers showed positive incoming MIF influence across low, intermediate and high levels of current melanocytic expression (**Fig. 4f**). The inferred circuit was thus predominantly tumour-associated but was not confined to a single expression state. Gene-level decomposition across the full signature further showed that programme decline emerged from the balance of marker responses, rather than uniform suppression of every melanocytic gene (**Supplementary Fig. 5d,e**).

The distinction between current state and predicted dynamics was apparent across sections. Incoming MIF influence correlated more strongly with predicted melanocytic decline than with current programme expression, and receivers in the upper quartile of positive incoming MIF influence had more negative total programme velocities than receivers with zero incoming MIF influence (**Fig. 4g**). These comparisons placed neighbourhood signalling alongside the direction of RNA change, identifying spatial contexts in which an established lineage programme was predicted to weaken without a correspondingly strong relationship to its observed abundance.

They also connected the local sender-to-gene decomposition to a predicted trend in melanocytic-programme change across malignant receiver spots while preserving the distinction between model-based attribution and experimentally measured effects. A plausible regulatory context is provided by experiments showing that MIF-CD74 blockade suppresses AKT signalling in melanoma^30^ and that AKT-dependent phosphorylation modulates MITF transcriptional function^31^ (**Extended Data Fig. 8a**). With measured RNA and fitted parameters held fixed, removing the MIF communication component attenuated predicted programme decline most clearly in S4, with a more modest response in S2 and little change in S3 (**Extended Data Fig. 8b,c**). Spatial projections and paired full-gene angles further showed directional responses, with the largest median angular change in S4 (**Extended Data Fig. 8d,e**). Independent MITF overexpression and knockdown produced opposing lineage-programme responses in A2058 cells (**GSE167496; Extended Data Fig. 8f**). Gene-resolved comparisons and increased shared-programme expression after *Mif* deletion supplied complementary expression-level evidence (**GSE247057; Extended Data Fig. 8g,h**). These independent perturbations contextualised the inferred response without establishing the complete MIF-AKT-MITF cascade in the spatial tissue.

### CellVocal Resolves Hepatocyte-Derived ANGPTL4 Signalling and Matrix-Remodelling RNA Dynamics in Acute Liver Failure

Tissue repair after acute liver failure requires local signals to be coordinated with hepatocyte transcriptional responses. We applied CellVocal to the human liver regeneration dataset reported by Matchett et al.^32^, comprising eight Visium sections from seven donors across healthy liver, acetaminophen-associated acute liver failure (APAP) and acute liver failure of non-A-to-E aetiology (NonAE). The original study identified ANXA2-positive migratory hepatocytes involved in necrotic wound closure, providing a setting in which to examine how spatial communication accompanies repair-associated RNA dynamics. We defined receivers by an inferred hepatocyte fraction of at least 0.50, retaining the capture spot as the observational unit, and mapped their cellular neighbourhoods across clinical conditions (**Fig. 5a**). This design allowed us to ask both which communication modules were preferentially deployed in APAP injury and how those modules contributed to the predicted responses of nearby hepatocytes. Negative binomial regression across 14 sender and ligand-receptor groups, with total event offsets and covariance clustered by donor, identified hepatocyte-derived ANGPTL4/SDC1 communication as enriched in APAP (**Fig. 5b**). The corresponding event rate was approximately 1.5-fold higher than in either healthy liver or NonAE (FDR < 0.001 for both contrasts), with greater receiver coverage and little difference between the two comparison groups (**Fig. 5c**). The enrichment was supported across the principal regression specifications (**Supplementary Fig. 6**), alongside a distinct APAP-associated organisation of donor communication profiles (**Extended Data Fig. 9a,b**). These comparisons positioned ANGPTL4/SDC1 as a candidate local signalling interaction associated with the APAP injury context rather than with acute liver failure irrespective of aetiology.

**Fig. 4.**
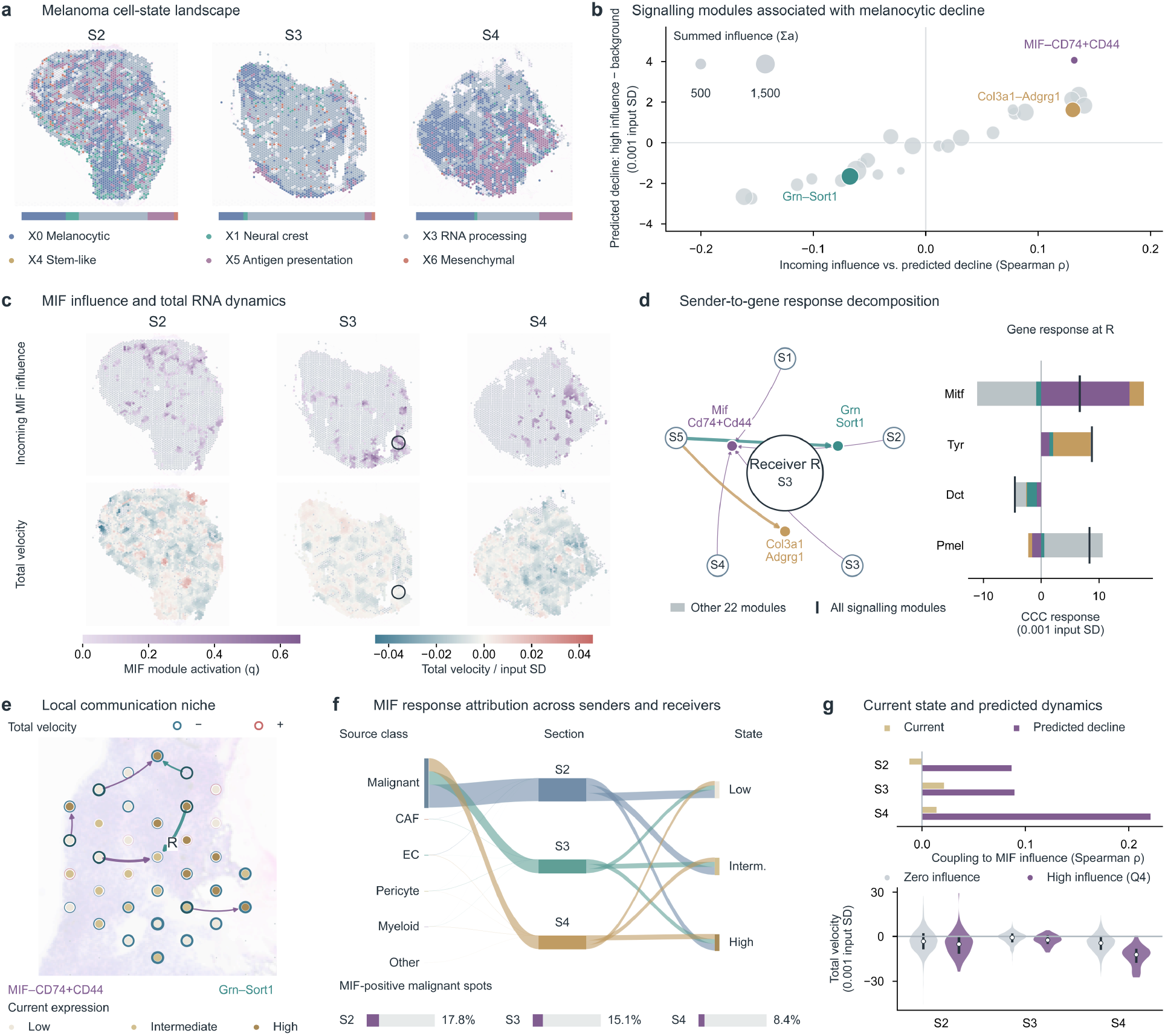
CellVocal resolves spatial MIF signalling associated with melanocytic RNA dynamics in melanoma. **a** Spatial distributions and proportions of malignant states across Visium sections S2-S4. **b** Screening of 25 ligand-receptor modules by section-averaged correlations between incoming signalling influence and predicted decline, together with contrasts between spots with high incoming influence and background spots. For each module, the high-influence group comprises spots in the upper quartile of positive incoming influence; background comprises remaining malignant spots. Bubble area represents summed signalling influence. Decline is negative total melanocytic-programme velocity. **c** Spatial distribution of incoming MIF-CD74+CD44 signalling influence and total programme velocity on shared scales. Circles mark the S3 neighbourhood in d,e. Gene velocities are scaled by input-expression standard deviations. **d** Inferred signalling influence of individual senders across three modules at receiver R and their contributions to *Mitf, Tyr, Dct* and *Pmel* communication-associated velocities. Grey bars combine the other 22 modules; black ticks mark the net response. **e** Local MIF-CD74+CD44 and GRN-SORT1 routes. Receiver fill indicates current expression, rings encode total velocity sign and magnitude, and arrow width represents attributed response magnitude. **f** MIF response attribution across sender classes, sections and receiver expression states, normalised by malignant-spot counts with equal section weighting. Bars show the fractions of receivers with positive incoming MIF influence. **g** Correlations of incoming MIF influence with current expression and predicted decline (top), and programme-velocity distributions in receivers with zero incoming MIF influence and those in the upper quartile of positive incoming MIF influence (bottom). Points and bars indicate medians and interquartile ranges.

**Fig. 5.**
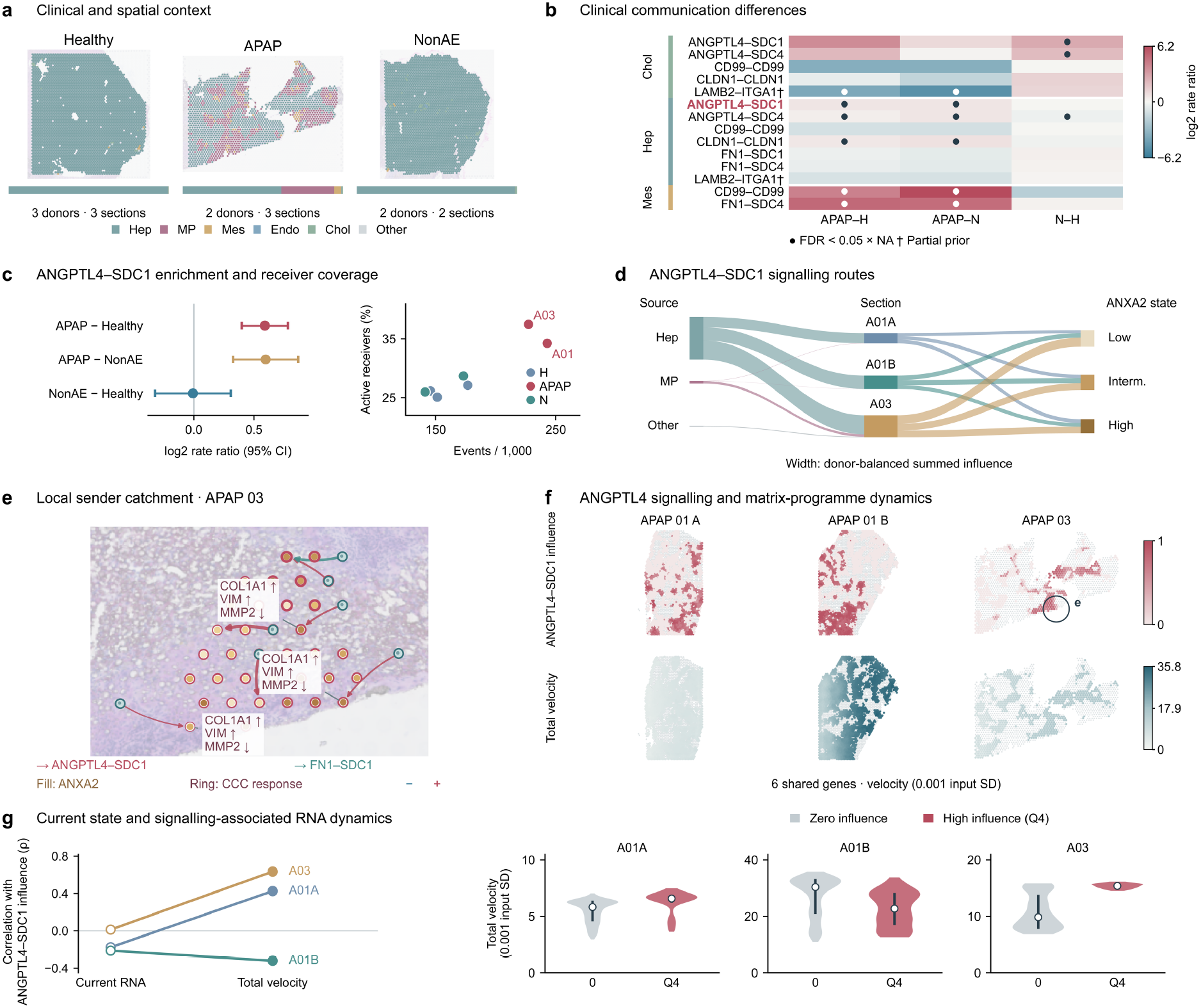
CellVocal resolves ANGPTL4 communication and associated RNA dynamics in acute liver failure. **a** Cell class distributions and proportions in representative healthy, APAP and NonAE sections. Eight sections represent seven donors; APAP01A and APAP01B share one donor. **b** Negative binomial contrasts across 14 sender and ligand-receptor groups, with total event offsets and covariance clustered by donor. Dots indicate FDR < 0.05 across 42 comparisons. **c** Hepatocyte-derived ANGPTL4/SDC1 contrasts with 95% confidence intervals, donor event frequencies and receiver coverage. **d** Signalling routes across sources, sections and ANXA2 expression states; ribbon widths show summed signalling influence, with each donor contributing equally. **e** Local sender catchment in APAP03. Fills show ANXA2 expression, rings encode signed communication-associated responses, and arrows indicate sender attribution. **f** Incoming ANGPTL4/SDC1 signalling influence and total RNA velocity for ANXA2, VIM, FN1, COL1A1, COL1A2 and ACTA2, with scales shared across sections within each row. The circle locates e. **g** Correlations of incoming ANGPTL4/SDC1 influence with current expression and total velocity, followed by velocity distributions for receivers with zero incoming ANGPTL4/SDC1 influence and those in the upper quartile of positive influence for this module. Points and bars indicate medians and interquartile ranges. Receivers have inferred hepatocyte fractions ≥0.5; velocities are scaled by gene-specific input standard deviations. †, incomplete receptor prior.

CellVocal then resolved how this condition-associated signalling interaction was distributed across sender populations, receiver states and individual neighbourhoods. With donor contributions balanced, hepatocyte-annotated senders accounted for most of the inferred ANGPTL4/SDC1 signalling influence, and inferred signalling reached receivers spanning low, intermediate and high ANXA2 expression states (**Fig. 5d**). The dominant inferred source therefore lay within the hepatocyte compartment itself, suggesting a local communication component accompanying the transcriptional response to injury. This attribution also separated the total incoming ANGPTL4/SDC1 influence from the identities of the contributing senders, a distinction that is not available from a receiver expression score alone. In an APAP03 neighbourhood, spatial sender catchments resolved ANGPTL4/SDC1 and FN1/SDC1 routes converging on individual receiver spots (**Fig. 5e**). Current ANXA2 expression, signed communication-associated responses, and gene-level directions could consequently be examined together without collapsing the neighbourhood into a single ligand-receptor score. The illustrated receivers combined positive COL1A1 and VIM responses with a negative MMP2 response, showing how communication-associated dynamics can distinguish components of a broader remodelling programme. Decomposition across the full set of modules extended this interpretation to overlapping signalling modules and their unequal contributions to receiver genes (**Extended Data Fig. 9c,d**). Such local organisation is biologically compatible with recent evidence that hepatocyte-neutrophil LIFR/STAT3 feedback and glutamate-induced macrophage WNT3 signalling regulate hepatic repair^33,34^. These established mechanisms provide context for interpreting spatially organised repair responses, while the ANGPTL4/SDC1 attribution identifies a candidate route for further investigation.

To relate these local response assignments to RNA dynamics across hepatocyte-enriched receiver spots, we compared incoming ANGPTL4/SDC1 influence with total RNA velocity for the six remodelling-associated genes shared by all three APAP sections: ANXA2, VIM, FN1, COL1A1, COL1A2 and ACTA2 (**Fig. 5f**). In APAP01A and APAP03, incoming ANGPTL4/SDC1 influence was positively associated with programme velocity despite weak or negative associations with current programme expression; the opposing association in APAP01B highlighted the influence of local tissue context (**Fig. 5g**). This comparison distinguished associations with the observed transcriptional state from associations with its predicted rate of change. We further removed ANGPTL4/SDC1 and ANGPTL4/SDC4 communication components separately and jointly while keeping RNA, latent states and fitted parameters fixed (**Extended Data Fig. 10a**). Within the model, joint removal of these communication components reduced mean programme velocity across all APAP sections, with spatially heterogeneous changes and negative mean shifts across the six genes (**Extended Data Fig. 10b-d**). Spatial projection differences showed the direction of the perturbation response, while paired full-gene angles remained small, with medians of 0.69-0.93° after joint removal (**Extended Data Fig. 10e,f**). Independently, author-released data from LIDPAD mouse liver showed lower six-gene expression after ANGPTL4 neutralisation or myeloid *Angptl4* deletion at eight weeks (https://github.com/chenghongsheng/Angptl4-in-NAFLD-immunopathologies; **Extended Data Fig. 10g**), consistent with the model programme-level response; gene-specific expression responses varied with sampling time (**Extended Data Fig. 10h**). The experimental comparison concerns endpoint RNA in a distinct injury setting, providing complementary support for ANGPTL4-associated programme regulation while leaving the receptor-specific mechanism in human APAP injury open.

## Discussion

Spatial transcriptomics is shifting tissue analysis from static descriptions of cellular composition towards the resolution of spatially resolved cell-state transitions. Yet RNA velocity and cell-cell communication have largely been treated as separate problems: the former describes where cells may be heading, whereas the latter infers what signals may be exchanged between neighbouring cells. CellVocal establishes a unified dynamical connection between the two by representing changes in receiver-cell states as the joint outcome of intrinsic and communication-induced transcriptional dynamics, and by further decomposing the extracellular component across signalling modules and individual senders. Results from simulations and diverse tissues, including the brain, gut and lung, show that this framework stably recovers directions of cell-state transitions within complex spatial structures while identifying candidate regulatory signals consistent with receiver-cell transcriptional dynamics. The principal contribution of CellVocal is therefore not simply to place RNA velocity and CCC analysis in the same pipeline, but to transform conventional communication networks based on co-expression and spatial proximity into networks of signalling influence constrained by inferred RNA velocity.

This design establishes a bidirectional interpretive relationship between RNA velocity and cell-cell communication. RNA velocity first constrains communication inference. The inferred direction of transcriptional change in a receiver cell provides an additional dynamical reference, assigning less influence to candidate events that are supported by ligand-receptor expression or spatial proximity but cannot account for the receiver’s subsequent state change. This constraint reduces the typically large candidate space produced by static CCC analysis. Communication, in turn, helps to interpret RNA velocity. CellVocal decomposes total RNA velocity into state-dependent intrinsic dynamics and a communication-induced component derived from signalling influence and module-specific gene effects, and subsequently traces the latter to spatial locations, signalling modules, sender cells and target genes. A velocity vector is therefore no longer restricted to describing the state towards which a cell is moving; it can also be interpreted in terms of the local extracellular signals that may contribute to the underlying transcriptional change. This relationship does not involve using velocity to validate communication and then using communication to validate velocity. Instead, expression, spatial context, temporal information and ligand-receptor priors provide complementary constraints within the same model: velocity supplies a directional criterion, whereas inferred signalling influence and module-specific gene effects together define the extracellular component of the velocity field.

This bidirectional relationship connects local signalling with predicted transcriptional change across biological settings. During embryonic neurogenesis, CellVocal linked neighbourhood NCAM1-NCAM1 signalling to lower cell-cycle-associated total RNA velocity at immature neuronal capture sites. In melanoma, local MIF-CD74+CD44 signalling corresponded to a predicted decline in the melanocytic programme. Analysis of human acute liver failure identified APAP-enriched hepatocyte-derived ANGPTL4-SDC1 communication and traced its distribution across local receivers and matrix-remodelling RNA responses. Spatial attribution resolved these patterns across neighbouring senders, signalling modules and target genes, while removal of communication components with model parameters fixed quantified their contributions to predicted total RNA velocity at unchanged observed states. The section-dependent associations in melanoma and liver further highlighted the importance of receiver state and local signalling context. Together, these analyses connect spatial cell-cell communication patterns to individual receivers’ inferred dynamics, supporting spatially specific hypotheses about how local signalling contributes to transcriptional change. CellVocal’s estimates of signalling influence link candidate regulatory signals to predicted transcriptional changes and should be interpreted in light of ligand-receptor priors, RNA velocity quality, spatial resolution and donor sampling.

## Methods

### Datasets

#### Synthetic benchmarks with controlled ground truth

The synthetic datasets were designed to evaluate the ability of CellVocal to recover developmental time, infer RNA velocity and identify cell-cell communication events against controlled ground truth. The datasets comprised three simulation panels covering developmental direction, signalling event attribution and differential communication, and included unspliced and spliced RNA layers, spatial coordinates, ground-truth latent time, a decomposition of spatial velocity and ground-truth communication events. The simulated developmental trajectories included linear, bilinear, radial and multiradial patterns. The signalling event attribution simulations implemented organiser-dominant, collective-redundant, contact-dependent, graded-paracrine and developmental-switch communication mechanisms. The differential-communication panel further included matched control and perturbed conditions for evaluating the detection of differential CCC events. Implementation details are provided in **Supplementary Note 3** and **Supplementary Table 1**.

#### Mouse cortical Slide-seq benchmark

The mouse brain cortex dataset comprised the Slide-seq spatial transcriptomic data reported by Gu et al.^9^ and was used to evaluate spatial RNA velocity and cell-cell communication inference by CellVocal in the layered cerebral cortex. The dataset contained spatial coordinates and spliced and unspliced expression layers. We used the spatial x coordinate as a proxy for developmental ordering and adopted the cortical developmental axis from the ventricular zone (VZ), through the subventricular and intermediate zones (SVZ/IZ), to the cortical plate (CP), together with regions such as layers 5 and 6 (L5/6), as biological references for evaluating the recovery of cell-state transitions across cortical layers and velocity directions. The dataset was also used to characterise cortical ligand-receptor communication strength, receiver-level module activity and sender-cell composition.

#### Embryonic mouse gut Stereo-seq benchmark

The mouse gut dataset comprised embryonic mouse gut Stereo-seq spatial transcriptomic data reported by Gu et al.^9^. The dataset exhibited a well-defined tissue architecture and contained major cellular states, including epithelial and mesenchymal states. We identified the tissue boundary using Delaunay triangulation and used the distance from each spot to its nearest boundary cell as a proxy for developmental position, representing an inside-out growth process. The dataset was used to evaluate developmental-direction inference by CellVocal within a complex tissue boundary and to characterise RNA velocity and the spatial communication patterns of representative LR pairs, including *Ncam1-Ncam1, Lamc1-Itgb1* and *Tenm3-Adgrl2*.

#### Embryonic mouse lung Stereo-seq benchmark

The mouse lung dataset comprised embryonic mouse lung Stereo-seq spatial transcriptomic data reported by Gu et al.^9^ and was used to test the generalisability of CellVocal in a branching developmental tissue. The dataset contained spatial coordinates and spliced and unspliced expression layers. As for the gut dataset, we identified the tissue boundary using Delaunay triangulation and used the distance from each spot to its nearest boundary cell as a proxy for developmental time, representing radial growth from the tissue interior towards the periphery. The dataset was used for the joint evaluation of latent-time inference, RNA velocity reconstruction and CCC-network inference, and for the analysis of spatial communication axes including *Mpzl1-Mpzl1, Slit2-Robo1* and *Ptn-Sdc2*.

#### SPTEdU-seq analysis of embryonic cortical neurogenesis

The mouse embryonic brain dataset was obtained from the SPTEdU-seq study by Niu et al.^26^. We selected one spatial transcriptomic section from each of three developmental stages: E12.5, E14.5 and E15.5. The UMI-filtered matrix provided by the authors was used as the total-expression layer. Spatial coordinates and UTI/EdU-related measurements were retained, including UTI count, UTI ratio, log1p-transformed UTI ratio and Gaussian mixture model (GMM) category. Starting from the raw sequencing reads, we generated spliced, unspliced and ambiguous counts using STARsolo in Velocyto mode and aligned these counts with the author-provided matrix by spatial barcode and gene symbol. Cell annotations were obtained from the Brain Knowledge Platform (https://brain-map.org/bkp/browse). After quality control, the dataset contained 20 region- or layer-resolved cell types, which were further consolidated into 10 major cell types. Immature inhibitory and excitatory neurons were designated as receiver cells for downstream communication analyses. Cell-cycle exit was defined as the negative within-stage z-score of the mean expression or RNA velocity of *Mki67* and *Top2a*. Separate scores were calculated from current expression and total RNA velocity; higher scores indicated lower current cell-cycle activity or a stronger predicted tendency towards cell-cycle exit, respectively. For each receiver barcode and ligand-receptor module, we summed the inferred signalling influence across candidate senders. Stratified analyses additionally used the log1p-transformed UTI ratio as an indicator of the current newborn-cell state.

#### Spatial analysis of mouse melanoma progression

The mouse melanoma dataset was obtained from the study by Karras et al.^29^ and comprised three 10x Genomics Visium sections from a primary melanoma model with the *Tyr*::CreERT2; *Braf*^V600E/+; *Pten*^fl/fl genotype. The original study provided the Visium spatial transcriptomic data and a matched single-cell reference atlas, and defined a hierarchy of melanoma cell states extending from proliferative-like to invasive or metastatic programmes. We supplemented the author-provided data with spliced and unspliced counts generated using velocyto. The single-cell reference matrix and its accompanying cell-type annotations were used as the reference for transferring major cell-type labels to Visium spots with Seurat. This procedure generated predicted labels and corresponding confidence scores for malignant cells, endothelial cells, cancer-associated fibroblasts (CAFs), pericytes and monocytes/macrophages, among other cell types. Malignant-state annotations (X0, X1, X3, X4, X5 and X6) were subsequently transferred to spatial spots using the malignant-cell reference annotations and were used to characterise the direction of malignant cell-state progression.

#### Spatial analysis of human liver regeneration

The human liver regeneration dataset was obtained from the study by Matchett et al.^32^ and comprised frozen human liver sections profiled using 10x Genomics Visium. The cohort covered healthy donors, acetaminophen-associated acute liver failure (APAP) and non-A-E acute liver failure (NonAE). Eight Visium sections were included in the analysis: three healthy sections, three APAP sections and two NonAE sections. Starting from the raw sequencing data provided by the authors, we generated spliced and unspliced expression layers using velocyto. To obtain spot-level functional annotations, we calculated module scores for predefined gene signatures from the normalised expression matrix. These signatures represented hepatocytes, myofibroblasts, portal zonation, central zonation, cycling cells and the *ANXA2*/migratory programme. The cycling score combined the S-phase and G2/M-phase cell-cycle scores, whereas the zonation score was defined as the standardised difference between the central and portal signature scores. For cell-composition analysis, we used the author-provided single-nucleus RNA-sequencing atlas as a reference and applied RCTD to decompose the expression profile of each Visium spot into proportions of the major cell types. This analysis generated spot-level cell-type proportion annotations for hepatocytes, mesenchymal cells and other major cellular populations.

### Model Formulation

Given a spatial transcriptomic section comprising *N* spatial locations and *G* genes, the inputs consist of an unspliced RNA count matrix *U*, a spliced RNA count matrix *S*, spatial coordinates *X* and a set of ligand-receptor priors ℳ, where 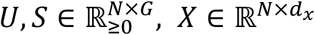. For a sender location *j*, a receiver location *i* and a communication module *m* ∈ ℳ within the spatial neighbourhood, a candidate communication event is defined as 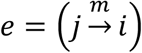.

CellVocal jointly infers the latent RNA state *z*_*i*_, relative kinetic time *τ*_*i*_, RNA velocity fields and the signalling influence 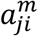 of candidate communication events,

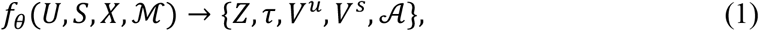

where 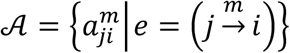 collects the influence estimates for all candidate sender-receiver-communication-module events. CellVocal first constructs a local adjacency graph from the spatial coordinates and generates candidate communication events between neighbouring locations. A graph neural network then integrates the local spatial context to learn denoised RNA states and latent representations, and estimates relative kinetic time and endogenous RNA velocity. Cell-type or tissue-region annotations can additionally be incorporated as optional priors. At its core, CellVocal decomposes the change in RNA state at location *i* into endogenous and communication-induced dynamics,

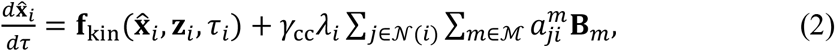

where the denoised RNA state is 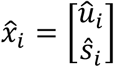 represents the endogenous dynamics arising from transcription, splicing and degradation; *B*_*m*_ is the gene-effect vector associated with communication module *m*; *λ*_*i*_ is the location-specific communication-response strength; and *γ*_cc_ is a global communication-scaling parameter.

For each receiver location, the model first evaluates the activities of different communication modules and subsequently estimates the relative share of each candidate sender within each module. Event-level signalling influence is factorised as 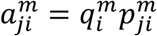, where 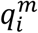 represents the activity of module *m* at receiver location *i*, and 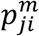 represents the relative contribution of sender location *j* within that module. Accordingly, the spliced RNA velocity at location *i* is expressed as

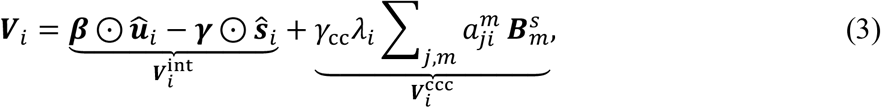

where 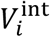 denotes the endogenous kinetic velocity and *V*^ccc^ denotes the exogenous velocity induced by cell-cell communication. CellVocal combines event-level influence estimates with module-specific gene-effect vectors to derive the communication-induced velocity, which, together with the endogenous dynamics, constitutes the total RNA velocity field. Thus, CellVocal is designed not merely to detect ligand-receptor co-expression between spatially proximal locations, but to map candidate communication events onto gene-level exogenous RNA velocity under the joint constraints of expression evidence, spatial neighbourhoods and RNA dynamics. It further quantifies the relative contribution of each event to the communication-induced velocity, thereby jointly recovering cell-state transitions, RNA velocity and directional cell-cell communication.

### RNA Preprocessing and Ligand-Receptor Prior Construction

CellVocal takes spliced and unspliced RNA counts and spatial coordinates from spatial transcriptomic data as inputs. To ensure robust RNA velocity estimation and spatial communication inference, we first performed quality control on both spatial locations and genes. Locations with insufficient RNA counts, inadequate detection of velocity-relevant genes or invalid spatial coordinates were removed. Genes were required to have sufficient shared counts in both the spliced and unspliced channels, after which the genes with the highest mean expression were retained for modelling.

By default, each location was required to contain at least 3 spliced counts, 3 unspliced counts and 3 detected genes; each gene was required to have at least 20 shared counts; and no more than 2,000 genes were retained. When sequencing-depth normalisation was applied, the same scaling factor, calculated from the combined total counts of the spliced and unspliced channels, was used for both channels. This procedure corrects for differences in sequencing depth while preserving the relative proportions of spliced and unspliced RNA. Ligand-receptor priors were obtained either from a user-provided interaction set or from CellChatDB. For multisubunit ligand or receptor complexes, an interaction was retained only when all resolvable subunits were present and detectable. The effective expression of complex *c* at spatial location *i*, denoted *E*_*ic*_, was defined as the minimum expression across all subunits in its subunit set 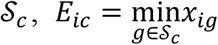, where *x*_*ig*_ denotes the expression of subunit gene *g* at spatial location *i*.

### Spatial Graph Construction and Candidate Communication Event Selection

#### Spatial neighbourhood graph construction

We constructed a symmetric *k* -nearest-neighbour graph *G*_sp_ = (*V*, ℰ_sp_) from the spatial coordinates to constrain local cell-cell communication. Self-loops were excluded. Candidate communication events were defined as directed events from sender location *j* to receiver location *i* within the spatial neighbourhood, where *j* ∈ N(*i*). The Euclidean distance *d*_*ji*_ associated with each edge was converted into a spatial decay weight,

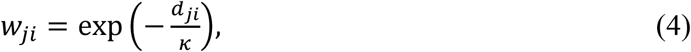

where *κ* was set by default to four times the median length of all edges with positive length.

#### Expression- and distance-aware candidate event selection

To mitigate the effect of zero inflation in spatial transcriptomic data on ligand-receptor detection, we first locally smoothed the spliced expression profiles over the spatial graph and then applied gene-wise min-max scaling to the smoothed expression values. For the *m*-th ligand-receptor module, the ligand expression *L*_*jm*_ at sender location *j* and the receptor expression *R*_*im*_ at receiver location *i* were both required to exceed the corresponding module-specific thresholds. These thresholds were determined jointly by a predefined lower bound and a global expression quantile, with the quantile set to 0.7 by default. For events passing the expression filters, the ligand and receptor expression levels were combined with spatial proximity to define a candidate-event ranking prior,

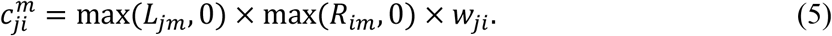

For each receiver-location-communication-module pair, only the *B* sender events with the highest 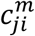 values were retained. This strategy limits the number of candidate events while preserving local competition among candidate senders. After candidate generation, communication modules containing fewer than *N*_min_ events were removed, with *N*_min_ = 10 by default. The remaining modules were then re-indexed for subsequent modelling.

### Graph-based RNA-State Representation and Latent-Time Inference

#### Input feature construction

To learn robust representations of kinetic state from sparse spatial transcriptomic observations, CellVocal jointly encodes the unspliced and spliced RNA signals at each spatial location and uses the spatial adjacency graph to integrate the local tissue microenvironment. Let *U, S* ∈ ℝ^*N*×*G*^ denote the unspliced and spliced count matrices, respectively, where *N* is the number of spatial locations, and *G* is the number of genes. We first applied a log(1 + *x*) transformation to *U* and *S* and concatenated the resulting features. When annotations such as cell type, cluster identity or tissue region were available, they were mapped to a learnable embedding *C* and supplied together with the RNA features as model input,

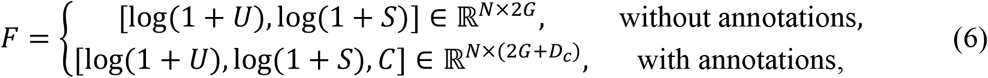

where 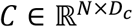 is the learnable embedding of the annotation information and *D*_*c*_ is its embedding dimension.

#### Graph-based latent RNA-state inference

CellVocal then uses a graph encoder *ϕ*_*θ*_ defined on the spatial graph *G*_sp_ to map the input features to a low-dimensional representation of RNA state,

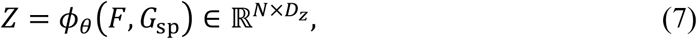

where 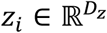 denotes the latent RNA state of spatial location *i*. A graph attention network is used as the encoder. By adaptively aggregating transcriptomic information from neighbouring locations, the estimated state of each location depends not only on its own spliced and unspliced signals but also on the transcriptional states of its local neighbourhood. This procedure reduces the effects of single-location measurement noise and zero inflation on velocity estimation.

#### Relative kinetic-time inference

Based on latent state *z*_*i*_, CellVocal further estimates the relative temporal ordering within a tissue section. Latent time is predicted by a nonlinear mapping *f*_time_ and constrained to the interval [0, 1] using a sigmoid function,

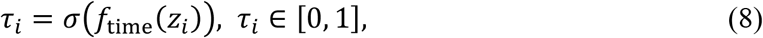

where *τ*_*i*_ represents the relative kinetic stage of spatial location *i* within the tissue rather than experimental time in absolute physical units. This representation provides a continuous state coordinate for subsequent RNA velocity estimation and enables velocity direction to remain consistent with local spatial transcriptional changes. In the implementation, *f*_time_ is a two-layer feed-forward network with a linear-ReLU-linear architecture.

### Context-Aware Representation of Communication Events

#### Multicomponent communication event encoding

For a candidate communication event 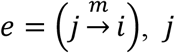 and *i* denote the spatial locations of the sender and receiver, respectively, and *m* denotes a communication module in the ligand-receptor prior library. Each module *m* corresponds to a set of ligand genes ℒ_*m*_ and receptor genes ℛ_*m*_. The event translator encodes event *e* as five complementary *d*-dimensional tokens,

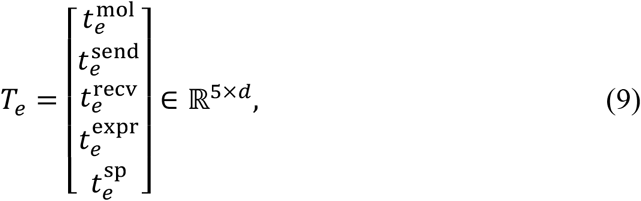

where 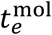 encodes the molecular identity of the communication module; 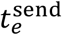 and 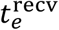 represent the local cellular states of the sender and receiver, respectively; 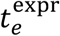 describes ligand-receptor expression compatibility; and 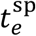 characterises the spatial relationship between the sender and receiver. Molecular identity is defined as 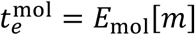, where *E*_mol_ is a learnable communication-module embedding layer. Each of the other four evidence components is mapped to the *d* -dimensional space through a component-specific learnable projection network 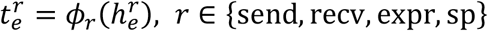. Their raw inputs are defined as

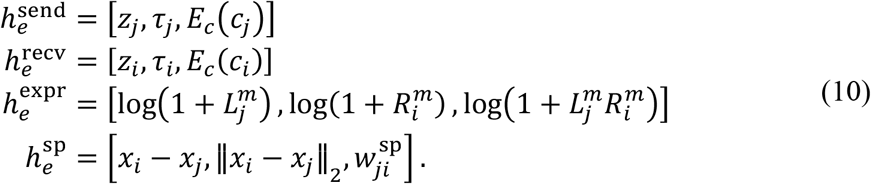

#### Query-based translation of event-level evidence

To extract event-level semantics from component-level evidence, the event translator first aggregates the event memory *T*_*e*_ into an event summary,

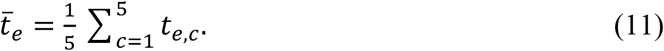

This summary represents the overall context of the candidate communication event and is used to modulate a set of learnable base soft queries. Specifically, a modulation network outputs scale and bias vectors 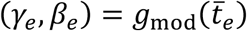. For the *r*-th base query 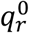, the event-specific query representation is defined as

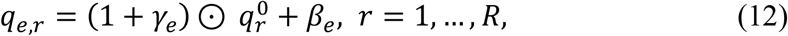

where *R* denotes the number of soft queries and ⊙ denotes element-wise multiplication. This modulation mechanism gives different candidate events distinct query states before they enter the attention layer, preventing all events from sharing an identical evidence-reading pattern. The event-specific queries then use *T*_*e*_ as keys and values in multi-head cross-attention,

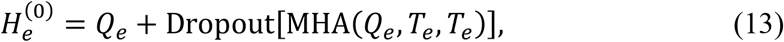

where *Q*_*e*_ = [*q*_*e*,1_, …, *q*_*e,R*_] ∈ ℝ^*R*×*d*^. This step enables multiple soft queries to retrieve complementary evidence in parallel from the same event memory, including communication-module identity, expression compatibility, spatial proximity and the state contexts of the sender and receiver. The cross-attended queries are subsequently passed through a Transformer encoder to promote information exchange among the soft queries and form higher-order event semantics,

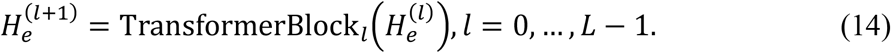

Finally, all soft queries are mean-pooled and subjected to layer normalisation to obtain an event-level semantic representation,

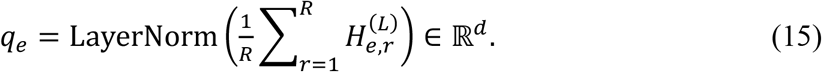

The resulting *q*_*e*_ provides a continuous semantic context for the candidate communication event and, after projection, is incorporated into the subsequent estimation of signalling influence. The event representation therefore does not depend solely on the presence of a particular ligand-receptor module, but jointly integrates module identity, sender state, receiver state, expression compatibility and spatial geometry.

#### Pretraining of the event translator

The event translator is pretrained independently before training the main model, such that its semantic event representations already contain stable structural information before entering the RNA-dynamics model. The pretraining objective comprises three components: communication-module classification, continuous-evidence regression and component-information reconstruction. First, event representation *q*_*e*_ is used to predict its communication module *m*_*e*_, thereby preserving categorical information about the ligand-receptor module,

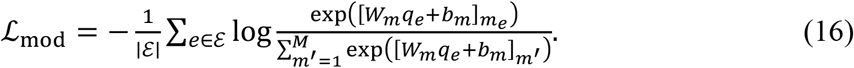

Second, to preserve continuous variation in the strength of communication evidence, the event translator regresses four scalar quantities from *q*_*e*_: ligand-receptor compatibility *ρ*_*e*_, the spatially weighted candidate-event score *c*_*e*_, ligand expression *L*_*e*_ and receptor expression *R*_*e*_ . The continuous-evidence regression head is defined as

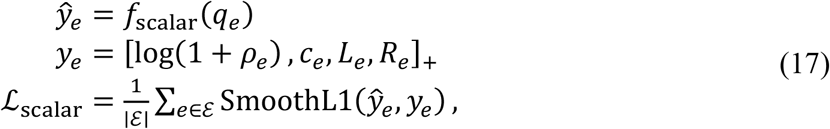

where *f*_scalar_ is a multilayer-perceptron regression head, 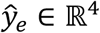 contains the predicted values of the four continuous communication features, and (·)_+_ denotes the element-wise non-negative part. This auxiliary constraint encourages the event semantic representation to preserve continuous information about ligand expression, receptor expression, ligand-receptor compatibility and the spatially weighted candidate score, rather than encoding only the discrete identity of the communication module. Finally, the translator is required to reconstruct the mean representation of its input components,

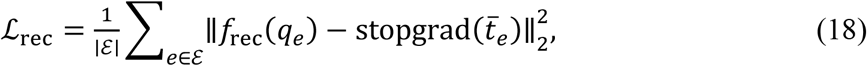

where stopgrad(·) makes the component-mean representation a stable reconstruction target and prevents the reconstruction branch from modifying the component encoders through backpropagation. The total pretraining loss of the event translator is defined as

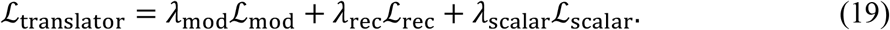

During pretraining, the receiver state is represented using a fixed state summary constructed from the spliced and unspliced RNA signals and does not depend on the latent representation subsequently learned by the main model. After pretraining, the translator parameters are frozen and a fixed semantic representation *q*_*e*_ is generated for every candidate communication event.

### Sparse inference of signalling influence

For a candidate communication event 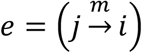, let 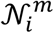 denote the set of candidate senders for communication module *m* at receiver location *i*. CellVocal factorises the inferred influence of this event into two complementary probabilistic components. The first is a receiver-side gate that characterises the extent to which module *m* is activated at location *i*. The second is the relative share attributed to each candidate sender 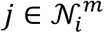, conditional on activation of module *m* . These two probability components are modelled using independent conditional categorical flows.

#### Conditional categorical-flow framework

For a probability vector *r*_*t*_ ∈ Δ^*K*−1^ with *K* states, the conditional categorical flow map is defined as

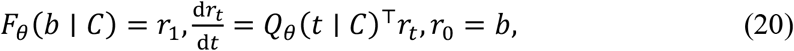

where *b* is the initial probability distribution, *C* is the conditional context, and *Q*_*θ*_(*t* ∣ *C*) is a neural-network-parameterised generator matrix for the categorical flow. Its off-diagonal elements are non-negative and the elements in each row sum to zero, ensuring that probability mass remains non-negative and sums to one throughout the flow. The resulting probability distribution can therefore be interpreted as the result of probability transport driven by the conditional context.

#### Estimation of conditional sender shares

For candidate sender location *j*, the sender-side communication representation is constructed as

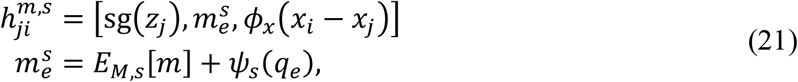

where *z*_*j*_ is the state of the sender location, *ϕ*_*x*_(*x*_*i*_ − *x*_*j*_) is the spatial-displacement encoding, *E*_*M,s*_[*m*] is the sender-side module embedding, *q*_*e*_ is the event semantic representation, and *ψ*_*s*_(·) is the sender-side semantic projection network. This representation jointly encodes sender state, event-module semantics and spatial propagation, but excludes receiver state *z*_*i*_ and ligand-receptor expression information, thereby maintaining independence between the sender-side prediction, receiver-side gate and expression prior. Candidate senders within the same receiver-module group are organised as an unordered set 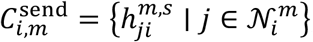, and a permutation-equivariant set network is used to parameterise the categorical flow. This construction ensures that the ordering of candidate senders does not affect their estimated shares and naturally accommodates candidate sets of different sizes. Using the uniform distribution 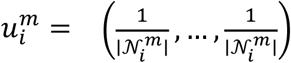 over candidate senders as its starting point, the conditional categorical flow yields the predicted vector of sender shares,

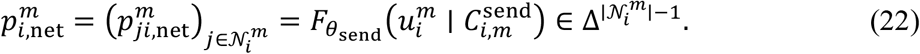

Uniform initialisation does not favour any sender a priori, allowing differences in sender shares to be jointly driven by sender state, module semantics and spatial relationships. Equation (22) transports probability mass jointly across the complete candidate set, thereby representing competition, substitution and cooperation among different senders.

#### Receiver-module activation inference

The receiver-side module gate evaluates whether module *m* is activated at receiver location *i*. Its conditional representation is defined as

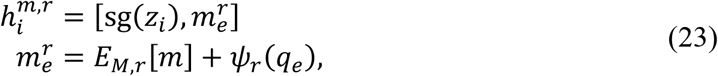

where *E*_*M,r*_[*m*] and *ψ*_*r*_(·) are the receiver-specific module embedding and semantic projection network, respectively, and their parameters are not shared with the sender-side branch. The receiver representation contains only receiver-location state and module semantics and is independent of the states, number and spatial distribution of candidate senders. The module state is represented by a binary distribution over inactive and active states. Starting from *u*_2_ = (1/2,1/2), the receiver-side network gate is obtained through an independent binary categorical flow,

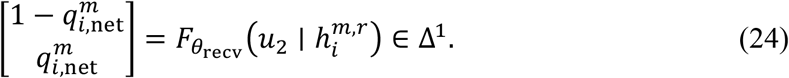

The first dimension represents the probability that the module is inactive, and the second represents the probability that it is active. This procedure interprets receiver-side gating as the transfer of probability mass between two module states and places it within the same categorical-flow framework as the estimation of sender shares.

#### Combining expression priors with learned probabilities

To retain the biological prior supplied by ligand-receptor expression, the prior for module activation and the prior for sender shares are constructed from receptor expression at the receiver and ligand expression at the sender, respectively,

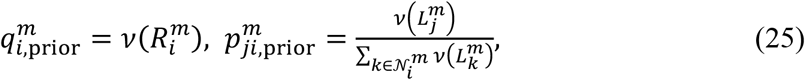

where 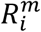 and 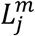 denote aggregated receptor expression at location *i* and aggregated ligand expression at location *j*, respectively. The robust expression-normalisation function *v*(·) compresses the expression dynamic range through non-negative truncation, a log(1 + *x*) transformation and a saturating map. The network branch therefore learns data-driven evidence supplied by latent state, event semantics and spatial structure, whereas the expression branch preserves an interpretable biological constraint. The final module activation probability and conditional sender shares are obtained by convexly combining the network predictions and expression priors,

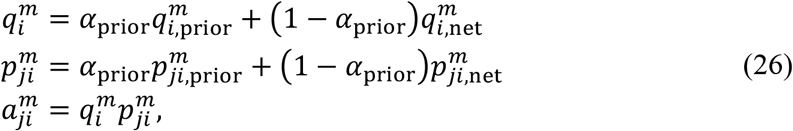

where *α*_prior_ ∈ [0, 1] controls the contribution of the expression prior. The quantity 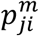 describes the relative contribution of candidate senders under a fixed receiver and module, whereas 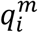 describes the overall activation of the module at the receiver location. Their product, 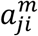, is the inferred signalling influence of candidate event *j* → *i* under module *m*.

### Decomposition of Endogenous and Communication-Induced RNA Velocity

For spatial location *i*, the model decomposes RNA velocity into an endogenous kinetic component and a communication-induced component. Given latent state *z*_*i*_ and latent time *τ*_*i*_, a state-modulation network first predicts gene-specific transcriptional modulation terms,

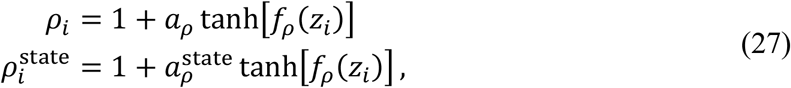

where 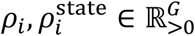 are used for velocity calculation and denoised-state reconstruction, respectively, and 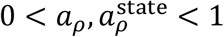 controls the amplitude of state-dependent modulation. The gene-level basal kinetic parameters 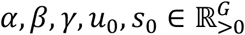 are parameterised using softplus to ensure non-negativity.

Given *τ*_*i*_, the model reconstructs the denoised RNA states 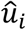 and 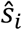 using the analytical solution of constant-coefficient splicing kinetics, with an effective transcription rate of 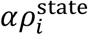.

Based on the denoised states, endogenous RNA velocity is defined as

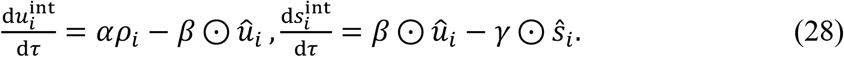

Communication-induced velocity is obtained by multiplying each event’s inferred influence by its module-specific gene-effect vector and summing across events. For each communication module *m*, the model learns two module-specific gene-effect vectors 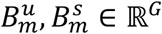, representing the exogenous effects of that module on the unspliced and spliced RNA derivatives, respectively. Given the event-level influence estimates 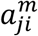, the communication-driven terms at receiver location *i* are

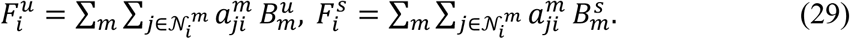

The model then predicts a location-specific communication-response strength from the denoised RNA state and latent state,

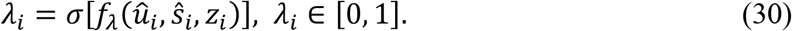

The communication-response gating network *f*_*λ*_ is implemented as a multilayer perceptron and mapped to [0, 1] using a sigmoid function, yielding the location-specific response strength *λ*_*i*_. The total RNA velocity field is then obtained by adding the endogenous kinetic velocity and the communication-induced velocity,

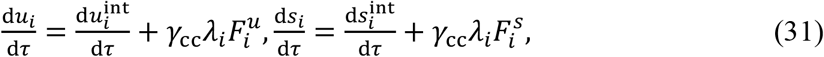

where *γ*_cc_ is the global communication-velocity scale. This additive decomposition enables the model to output the endogenous velocity, communication-induced velocity and combined RNA velocity separately, while tracing communication effects to spatial locations, communication modules and target genes.

### Training Objective

CellVocal jointly optimises RNA dynamics, latent time and signalling influence through three complementary objectives,

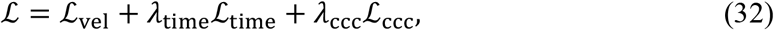

where ℒ_vel_ constrains the predicted velocity to agree with local transcriptional changes, ℒ_time_ constrains latent time to agree with an external or data-driven temporal prior, and ℒ_ccc_ aligns estimates of signalling influence, sender shares and module activation with ligand-receptor evidence for spatially defined candidates. This joint objective allows CellVocal to learn velocity direction directly from state changes within spatial neighbourhoods without requiring precomputed RNA velocity labels.

#### RNA velocity consistency objective

The model constructs supervision for RNA velocity from local expression changes on the spatial adjacency graph. For each valid adjacent edge, the earlier and later endpoints are first determined from a shared temporal prior, and only local relationships with a temporal difference exceeding a specified threshold are retained. For multiple later neighbours of the same earlier location, local least-squares regression is used to estimate the rates of change in unspliced and spliced RNA with respect to the temporal prior, yielding target velocities 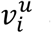 and 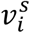. To prevent high-variance genes from dominating training, both predicted and target velocities are standardised by gene scale before comparison. A single-channel velocity-matching function is defined as

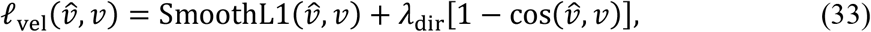

where 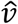 and *v* denote the standardised predicted and target velocities, respectively. The final RNA velocity consistency loss is

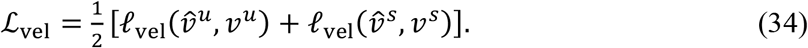

This loss constrains both velocity magnitude and direction, preventing the model from fitting only the magnitude of local expression changes while ignoring the direction of RNA-state progression.

#### Latent-time ordering objective

The direction of latent time is constrained by a dataset-specific temporal prior *t*^0^. This prior can be constructed from RNA maturity, a spatial coordinate or another continuous variable. By default, RNA maturity is defined as the difference between mean spliced expression and mean unspliced expression. The prior can be further smoothed over the spatial graph and standardised to zero mean and unit variance. The model randomly samples location pairs (*a, b*) with sufficiently different prior values and applies a pairwise ranking constraint to the pre-sigmoid time scores *h*_*i*_:

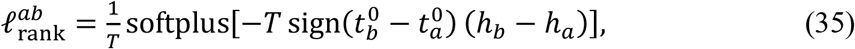

where *T* is a temperature parameter. The latent-time loss is

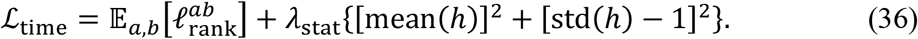

The first term aligns the predicted temporal ordering with the direction of the prior, whereas the second prevents the temporal scores from collapsing to a constant, thereby providing a continuous state coordinate with sufficient dynamic range for RNA velocity estimation.

#### Communication-prior consistency objective

Weakly supervised expression priors guide estimates of signalling influence, sender shares and module activation while allowing the RNA-dynamics objective to refine them. We define a within-group normalisation operator as

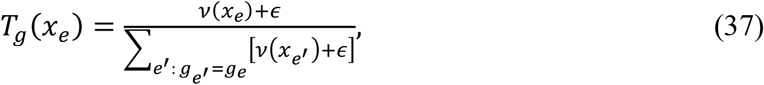

where *v*(·) is a normalisation function that robustly maps non-negative evidence to [0, 1], and *g*_*e*_ denotes the normalisation group. The communication-prior consistency loss is defined as

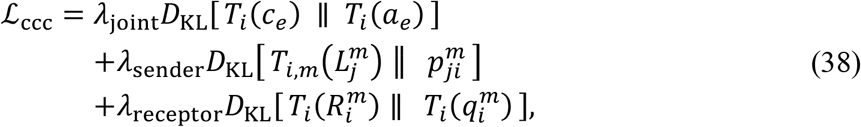

where 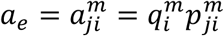 is the event-level estimate of signalling influence and *c*_*e*_ is the candidate-event score. The first term constrains the distribution of inferred influence across candidate events at each receiver location to agree with the candidate communication evidence. The second term constrains sender shares within each receiver-module group to agree with sender-side ligand expression. The third term constrains module activation within each receiver location to agree with receiver-side receptor expression. This loss preserves interpretable ligand-receptor priors during the early stages of training while allowing the RNA velocity consistency objective to refine the estimates of signalling influence.

## Data Availability

All datasets analysed in this study are publicly available from the repositories associated with the original publications. The human liver regeneration dataset is available through the Gene Expression Omnibus (GEO) under accession code **GSE223561**, and the mouse melanoma dataset is available under accession code **GSE207592**. Raw sequencing data from SPTEdU-seq are available under accession code **GSE283159**; the corresponding processed expression matrices and annotations are available from Figshare at https://doi.org/10.6084/m9.figshare.27923973. The mouse cortical Slide-seq dataset and the mouse gut and lung Stereo-seq benchmark datasets were obtained from the preprocessed AnnData datasets provided with TopoVelo (Figshare: https://doi.org/10.6084/m9.figshare.28516139.v2). Independent perturbation data used in Extended Data Figs. 6, 8 and 10 were obtained from GEO (**GSE228315, GSE167496** and **GSE247057**) and the authors’ public GitHub Repository . The ligand-receptor prior was obtained from CellChatDB.

## Code Availability

Source code for the CellVocal Python toolkit and scripts used to reproduce the analyses is publicly available at https://github.com/xuyungang/CellVocal.

## Acknowledgements

This work was supported by the National Natural Science Foundation of China (62471378, 82541006, 62401452, and 62501463), the Shaanxi Provincial Natural Science Foundation (No. 2024SF-GJHX-40, 2026JC-JCQN144), the Science Foundation of Xi’an Jiaotong University (No.YXJLRH2022011), and the Young Talent Support Plan of Xi’an Jiaotong University (No. YX6J021).

## Author Contributions

S.W. designed the study, performed the experiments, analysed and interpreted the data, and wrote and revised the manuscript. J.W. (Jiayin Wang) contributed to the study design. X.L. and J.W. (Jiayi Wang) contributed to manuscript revision. Y.X. designed the study, revised the manuscript, and acquired funding. All authors reviewed and approved the final manuscript.

## Extended Data Figures

**Extended Data Fig. 1.**
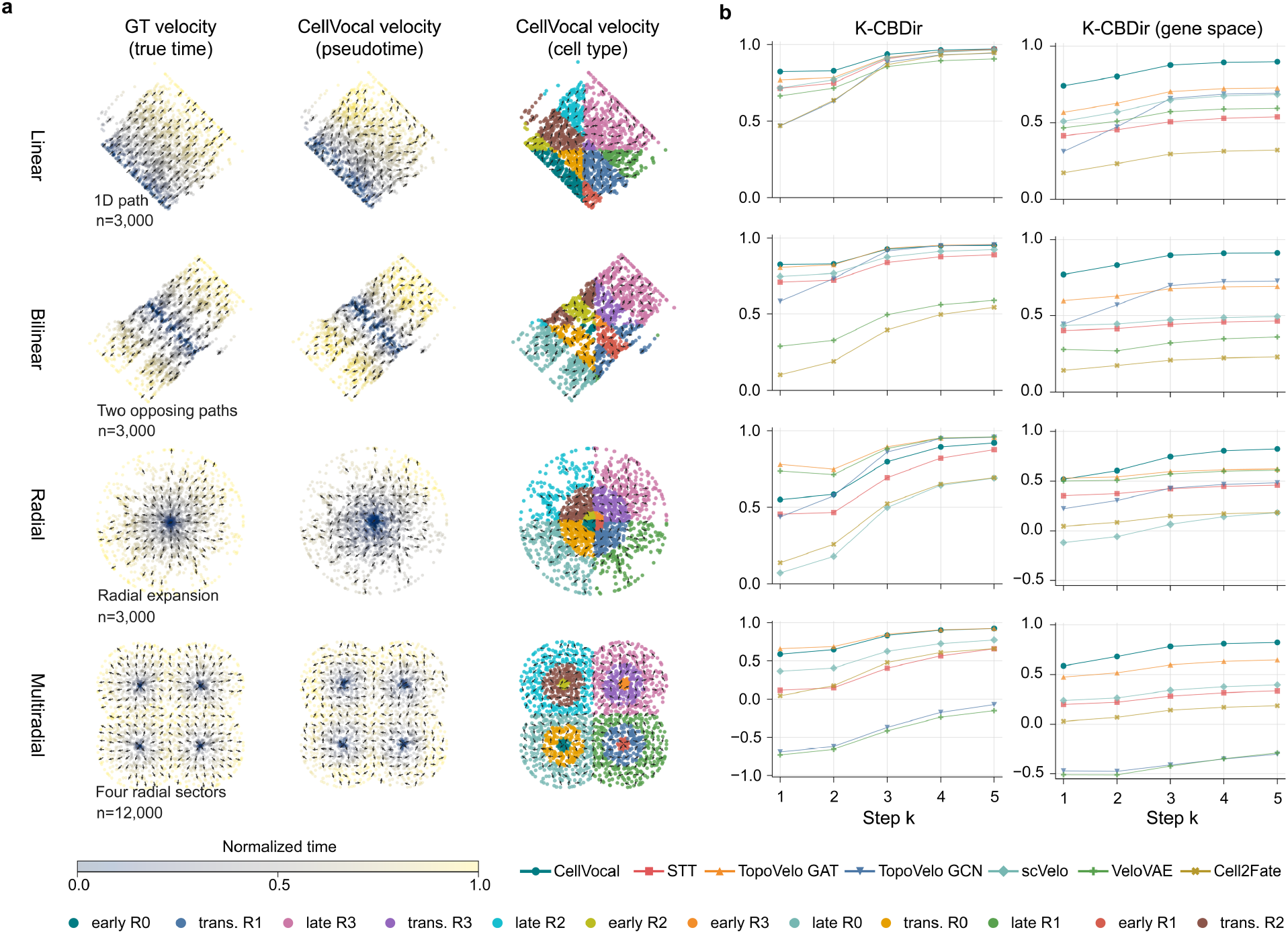
Directional velocity recovery across simulated spatial trajectories. **a** Ground-truth velocity fields and CellVocal-inferred velocity fields are shown for four simulated spatial trajectories: linear (1D path, *n* = 3,000), bilinear (two opposing paths, *n* = 3,000), radial (radial expansion, *n* = 3,000) and multiradial (four radial sectors, *n* = 12,000). The first column uses the known simulation time to define the ground-truth velocity, whereas the second and third columns show CellVocal velocity projected onto inferred pseudotime and simulated cell-type labels, respectively. Points are coloured by normalised time or cell type, and arrows represent local mean velocity directions. **b** K-CBDir and gene-space K-CBDir are plotted across prediction steps *k* = 1-5 for CellVocal and comparator methods. Each curve follows the same simulation topology across increasing prediction horizons; higher values indicate more accurate recovery of the underlying transition direction.

**Extended Data Fig. 2.**
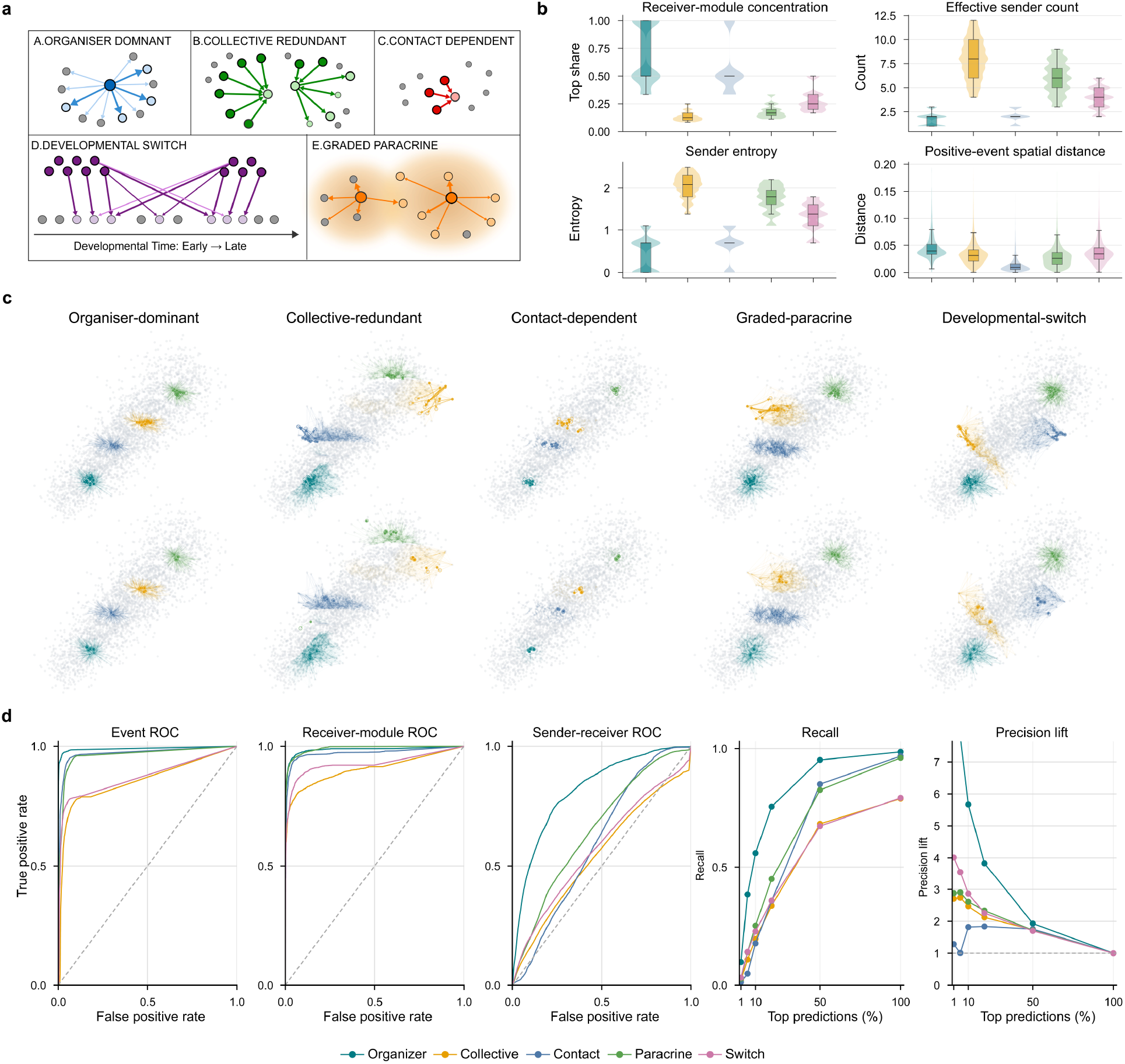
Recovery of simulated cell-cell communication mechanisms. **a** Schematic representations of five synthetic CCC mechanisms: organiser-dominant, collective-redundant, contact-dependent, graded-paracrine and developmental-switch communication. Nodes represent cells, arrows represent directed sender-to-receiver interactions, and coloured structures indicate the defining mechanism. **b** Ground-truth communication complexity quantified at the receiver-module level by top-sender share, effective sender count and sender entropy, together with the spatial distance of positive CCC events. Distributions are shown across receiver-module groups for each mechanism. **c** Spatial CCC networks in which the upper row shows ground-truth interactions and the lower row shows CellVocal predictions. Cells are displayed as a spatial background, while coloured arrows identify LR-specific directed edges. **d** Recovery performance evaluated at three resolutions—individual events, receiver modules and sender-receiver pairs using ROC curves. Recall and precision lift are additionally shown as a function of the fraction of top-ranked predictions retained. Colours identify the five simulated mechanisms.

**Extended Data Fig. 3.**
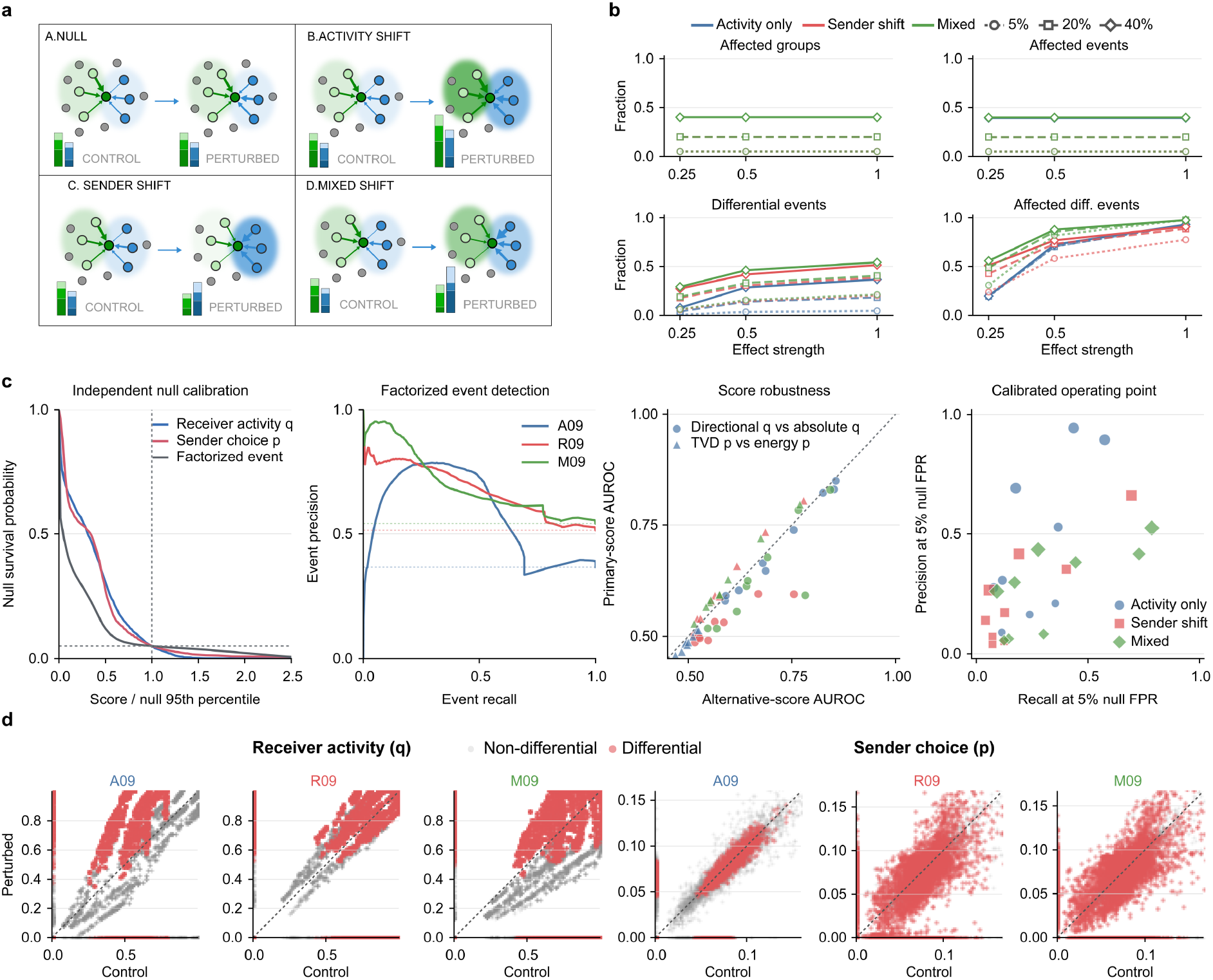
Comprehensive validation of differential CCC recovery. **a** Schematic of null, receiver-activity, sender-shift and mixed perturbations. Activity-only perturbations alter receiver activity while preserving sender composition; sender-shift perturbations alter the dominant sender while preserving receiver activity; mixed perturbations change both components. **b** Fractions of affected receiver groups, affected events, differential events and affected differential events across effect strengths of 0.25, 0.5 and 1.0 and affected fractions of 5%, 20% and 40%. **c** Validation of differential-CCC detection. From left to right: survival distributions of null-normalised scores for receiver activity (q), sender choice (p) and factorised events, with the null 95th-percentile threshold indicated; precision-recall curves for factorised event detection in representative activity-only (A09), sender-shift (R09) and mixed (M09) experiments, with dashed lines indicating event prevalence; robustness of primary AUROC values to alternative definitions of the receiver-activity and sender-choice scores; and precision versus recall at operating points calibrated to a 5% null false-positive rate. **d** Control-versus-perturbed scores for receiver activity (q) and sender choice (p) for events in A09, R09 and M09. Grey and red points denote non-differential and differential events, respectively; dashed diagonals indicate equal scores between conditions.

**Extended Data Fig. 4.**
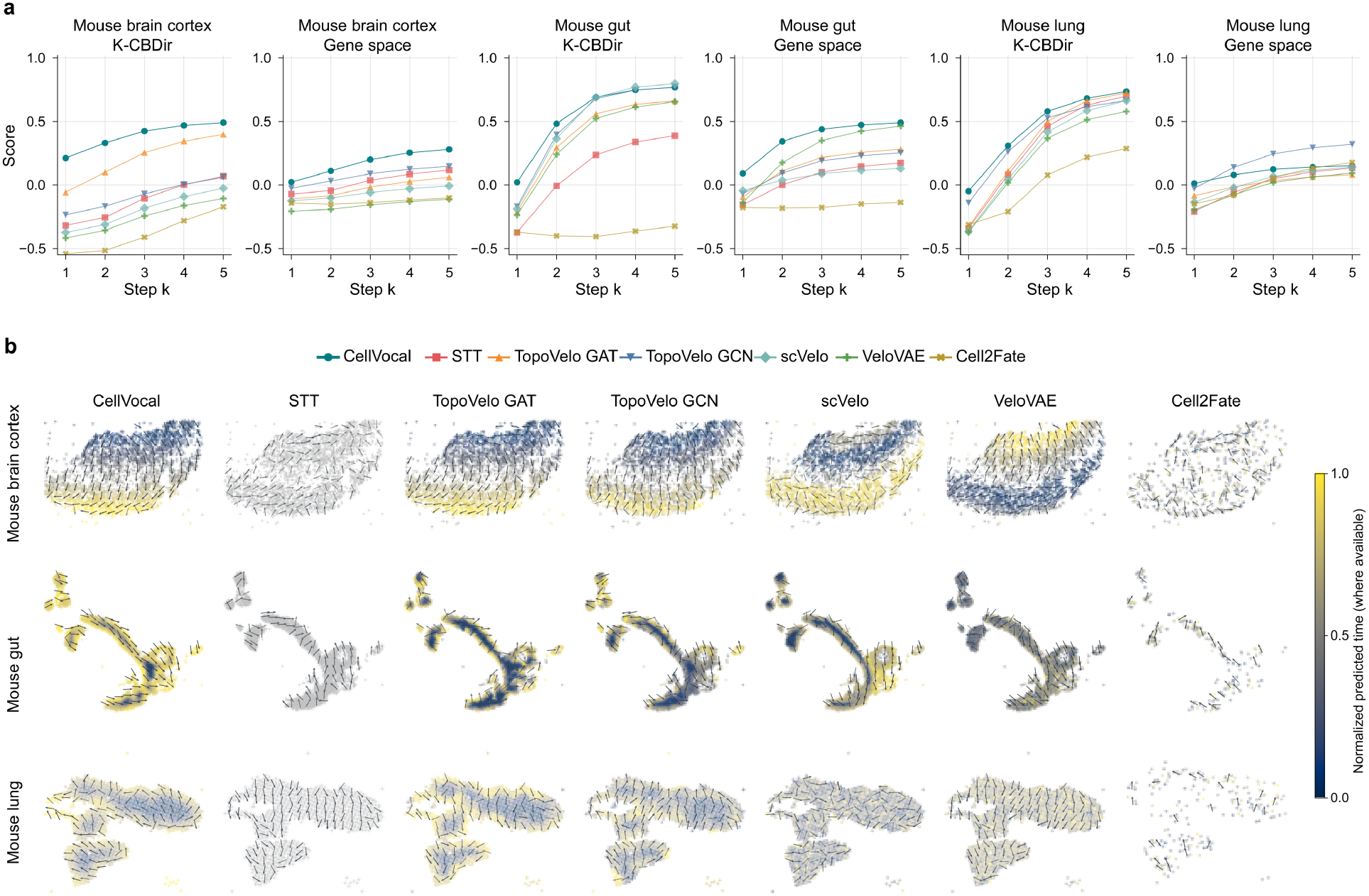
RNA velocity benchmarks across methods on real spatial transcriptomic data. **a** K-CBDir and gene-space K-CBDir curves across prediction steps *k* = 1-5 in mouse brain cortex, gut and lung for CellVocal and comparator methods. The curves assess how well each method preserves local transition direction over progressively longer prediction steps. **b** Spatial velocity fields inferred by CellVocal, STT, TopoVelo, scVelo, VeloVAE and Cell2fate in the same three tissues. The figure is arranged as a tissue-by-method grid, enabling direct comparison of the inferred spatial flow patterns. Points are coloured by normalised latent time when a method provides a time estimate; STT cells are shown in grey because no corresponding latent-time estimate is available. Arrows indicate locally coherent velocity directions, and CellVocal is highlighted in teal.

**Extended Data Fig. 5.**
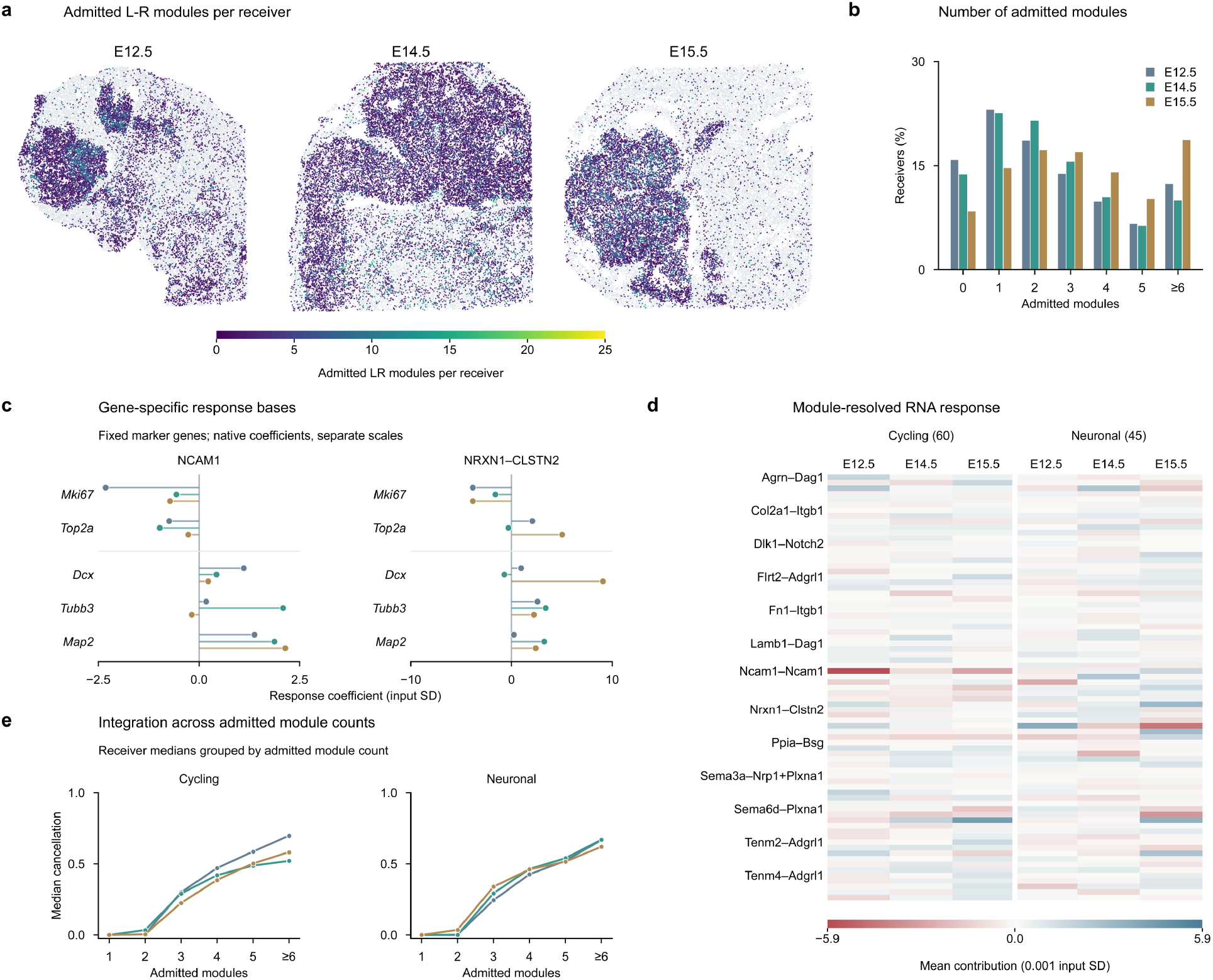
Decomposition and integration of multiple signalling contributions by CellVocal. **a** Spatial distribution of admitted ligand-receptor modules, defined by positive incoming signalling influence at the receiver, at immature neuronal receiver sites across three stages. **b** Receiver percentages by admitted-module count. **c** Signed gene-response coefficients for NCAM1-NCAM1 and NRXN1-CLSTN2 across five cycling and neuronal markers, scaled by gene-specific input standard deviations. **d** Mean contributions of all 77 modules to the 60-gene cycling and 45-gene neuronal-function programmes, incorporating incoming signalling influence, response modulation and gene-response coefficients. **e** Median cancellation of opposing module contributions by the number of admitted modules. Cancellation is calculated as 1 − |∑_*m*_ *c*_*im*_|/ ∑_*m*_|*c*_*im*_|, where *c*_*im*_ denotes the contribution of module *m* to receiver *i* for the indicated programme; larger values indicate stronger cancellation. Receivers with zero summed absolute contribution are excluded.

**Extended Data Fig. 6.**
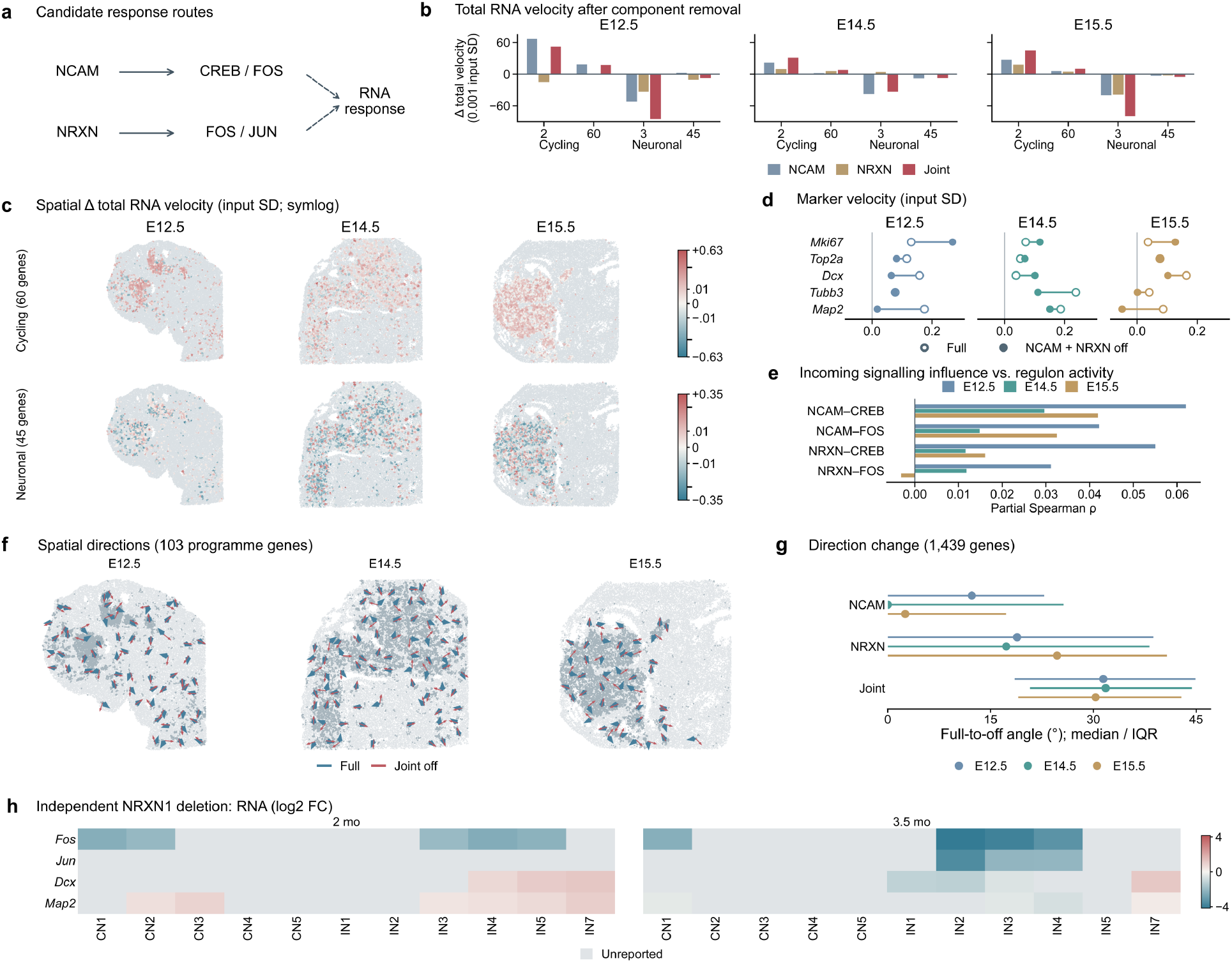
Neuronal RNA-velocity responses to in silico removal of NCAM and NRXN communication components. **a** Candidate adhesion-response routes. **b** Mean total RNA-velocity changes after separate or joint in silico removal of NCAM and NRXN communication components at E12.5, E14.5 and E15.5, evaluated at the same receivers with positive incoming influence from the targeted modules within each section. Cycling and neuronal readouts comprise marker sets of two and three genes and broader programmes of 60 and 45 genes. Model parameters, measured RNA and latent states remain fixed. **c** Spatial distributions of joint-component-off minus full-model programme velocity; symmetric logarithmic scales are shared across stages within each row. **d** Mean velocities of *Mki67, Top2a, Dcx, Tubb3* and *Map2* under full and joint-component-off conditions, shown by connected open and filled circles. Velocities in b-d are scaled by gene-specific input standard deviations. **e** Partial Spearman correlations between incoming signalling influence and CREB/FOS regulon scores, adjusted for spatial location, RNA depth, receiver lineage and current marker expression. Regulons exclude outcome and ligand-receptor genes. **f** Local spatial projections of full-model (blue) and joint-component-off (red) RNA velocity using the 103-gene union of the cycling and neuronal programmes. Arrows show spatially binned unit directions on the original tissue coordinates. **g** Paired full-to-component-off angles across 1,439 shared genes after scaling by gene-specific input standard deviations; points and horizontal lines indicate within-section medians and interquartile ranges. **h** Published expression contrasts after isogenic NRXN1 deletion in human brain organoids at two and 3.5 months. Heatmaps show reported log2 fold changes by neuronal class; grey indicates unreported values.

**Extended Data Fig. 7.**
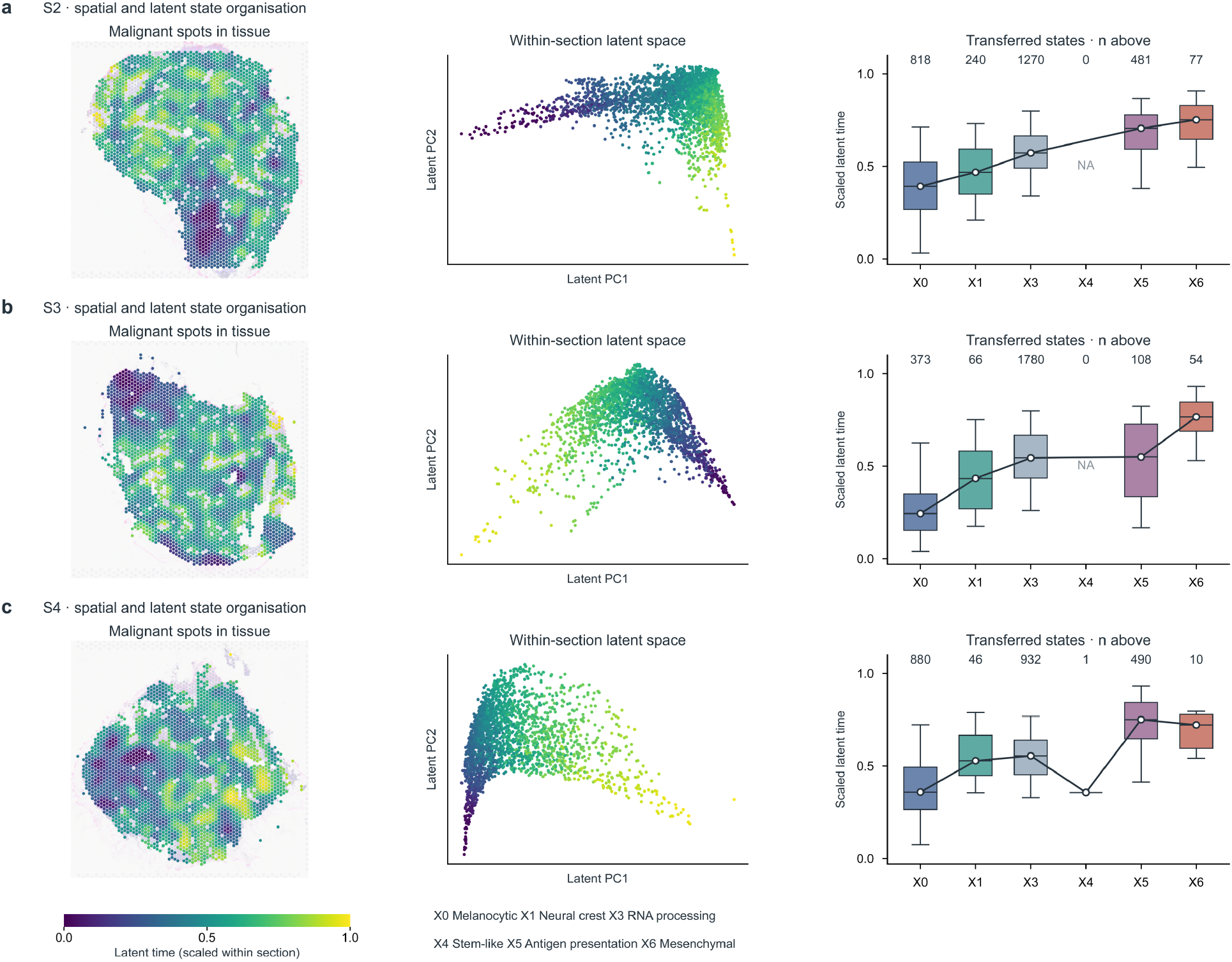
Spatial and latent organisation of melanoma cell states inferred by CellVocal. **a-c** Malignant capture spots in S2 (a), S3 (b) and S4 (c), shown in tissue coordinates (left), principal-component projections of inferred latent representations (middle), and latent-time distributions across transferred states (right). Map colours indicate latent time scaled to 0-1 within each section. Boxes span interquartile ranges, horizontal lines mark medians, and whiskers show the 5th-95th percentiles. Connected white points indicate state medians; numbers denote spot counts, and NA marks absent states. Transferred state order informed training, so these views describe the fitted spatial and state organisation.

**Extended Data Fig. 8.**
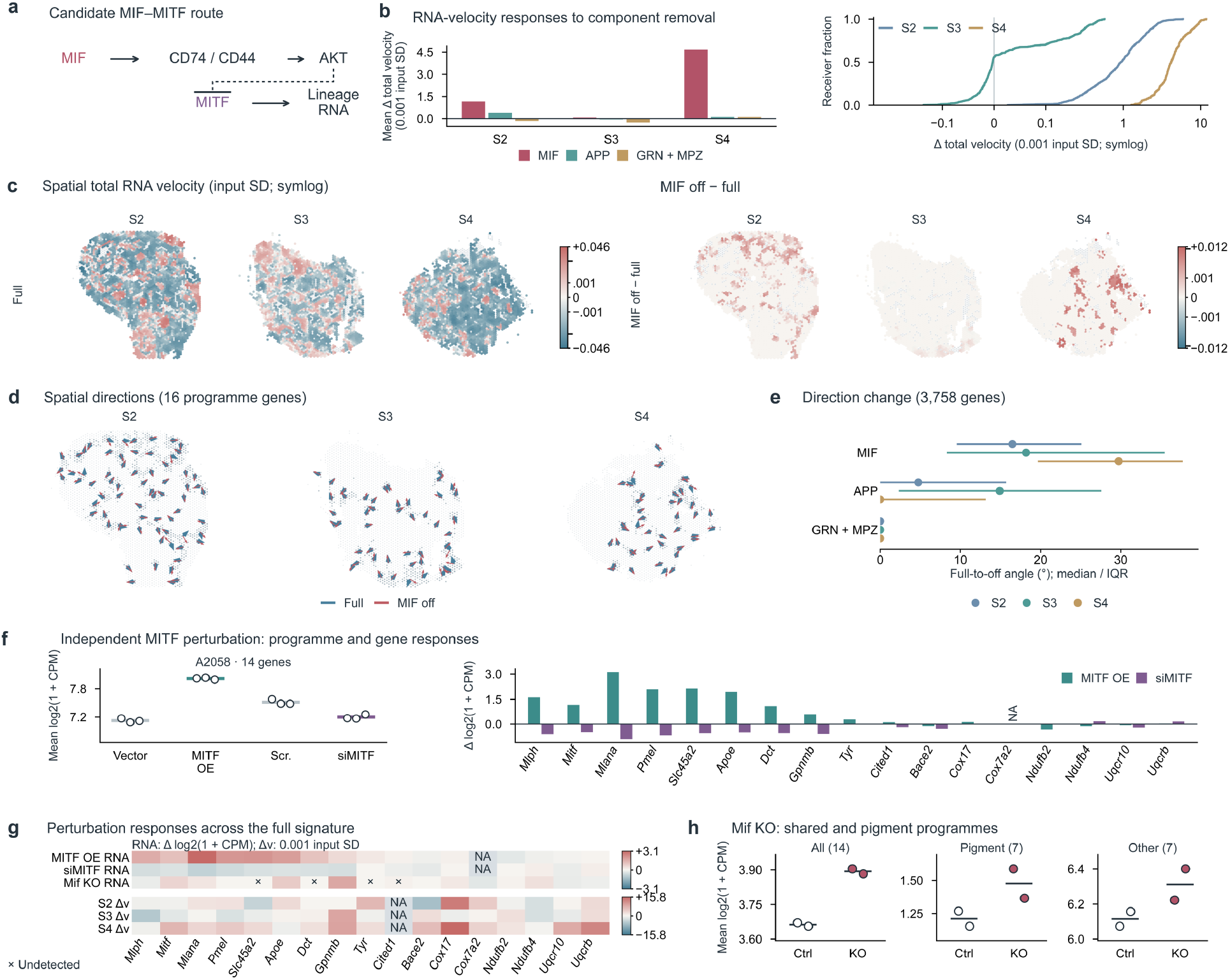
Dependence of melanocytic RNA dynamics on inferred MIF signalling and independent perturbation responses. **a** Candidate MIF-CD74/CD44, AKT and MITF response route. **b** Mean programme-velocity changes after in silico removal of MIF, APP or GRN + MPZ communication components in the same MIF-positive receiver cohort (left), and cumulative receiver distributions for MIF removal (right). Comparison signalling modules have different receiver coverage. **c** Spatial full-model total programme velocity (left) and MIF-component-off minus full-model changes (right) across S2-S4, with parameters, measured RNA and latent states fixed. Velocities in b,c are scaled by gene-specific input standard deviations; symmetric logarithmic colour scales in c are shared across sections within each group. **d** Local spatial projections of full-model (blue) and MIF-component-off (red) RNA velocity using 16 measured melanocytic-signature genes. Arrows show spatially binned unit directions on the original tissue coordinates. **e** Paired full-to-component-off angles across 3,758 shared genes after scaling by gene-specific input standard deviations; points and horizontal lines indicate within-section medians and interquartile ranges. **f** Independent MITF overexpression (OE) and knockdown in A2058 melanoma cells. Left, mean log2(1 + CPM) across 14 shared signature genes, excluding MITF; points show three biological replicates per condition, with means ± s.e.m. Right, gene-level changes relative to corresponding controls. **g** Expression contrasts after MITF OE, MITF knockdown and *Mif* knockout (top), and predicted MIF-component-off velocity changes (bottom), across all 17 signature genes. RNA and velocity use separate colour scales; NA denotes unmeasured or unmapped genes, and crosses mark undetected expression in *Mif*-knockout controls. **h** Expression of the shared 14-gene programme, seven pigment-associated markers and remaining seven genes in independent day-0 SCC35 melanoma cultures after *Mif* knockout. Points represent two independent clones per condition; horizontal lines show means.

**Extended Data Fig. 9.**
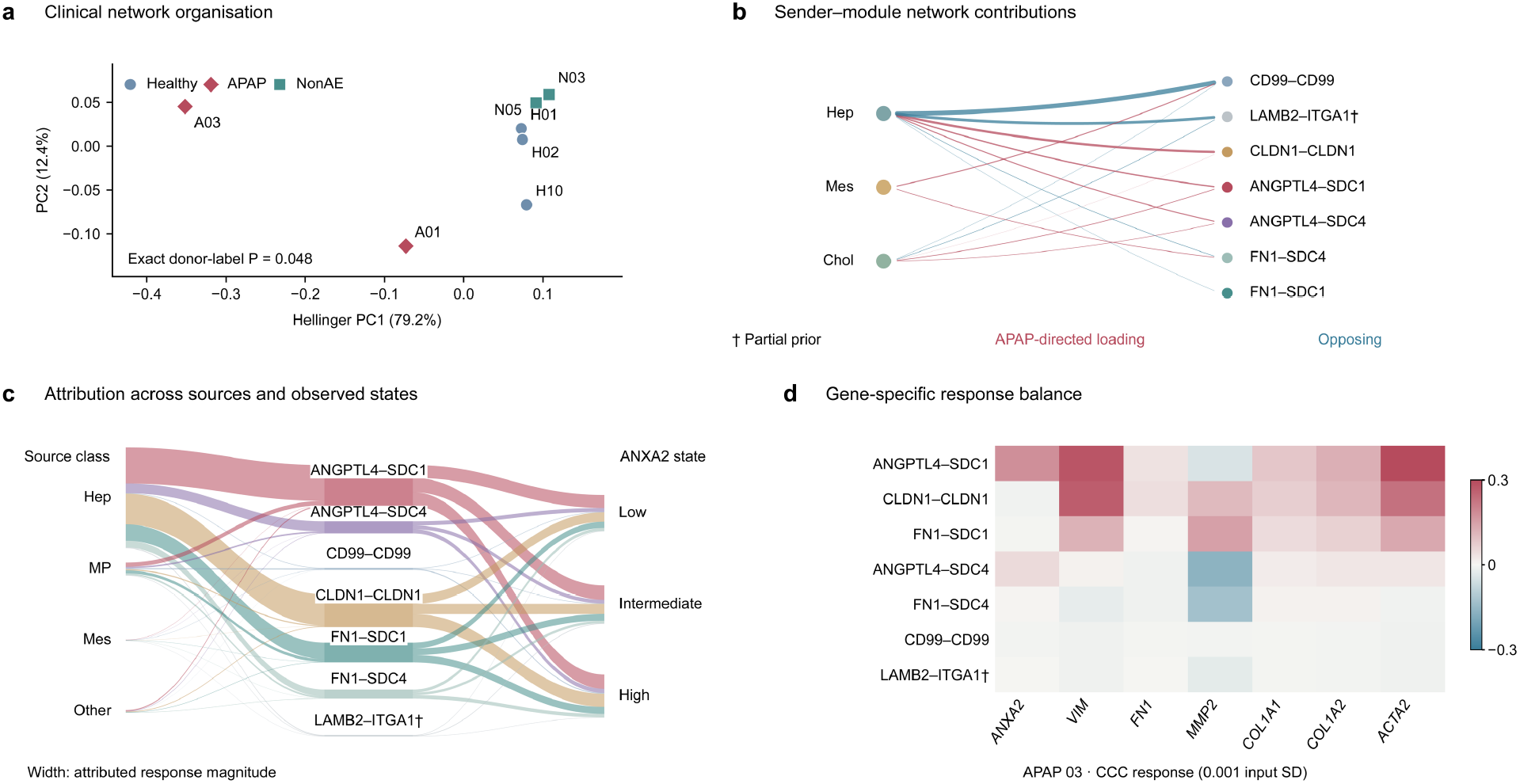
Decomposition of hepatic communication networks across donors, senders and gene responses. **a** Principal component analysis of Hellinger-transformed communication profiles from seven donors. The pseudo-F test uses all 21 APAP donor assignments. **b** Sender and module contributions through PC1 loadings oriented towards APAP. Edge widths indicate absolute loadings; colours distinguish APAP-directed and opposing contributions. **c** Attribution of response magnitude across sender classes, modules and ANXA2 expression tertiles in APAP03. **d** Mean signed contributions of seven modules to seven gene velocities, scaled by gene-specific input standard deviations. †, incomplete receptor prior.

**Extended Data Fig. 10.**
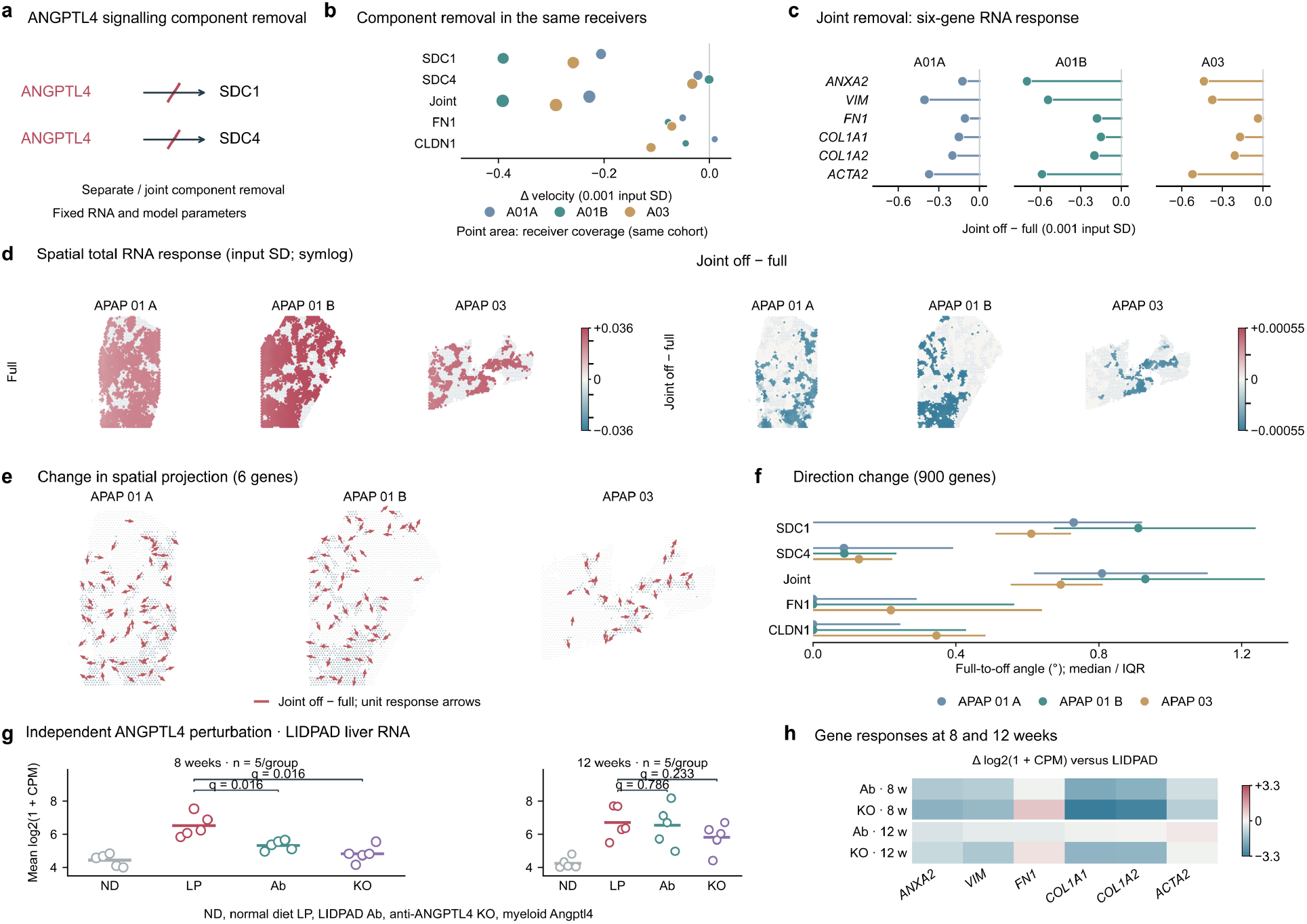
Dependence of predicted RNA dynamics on inferred ANGPTL4 signalling and independent liver perturbation responses. **a** Separate and joint removal of ANGPTL4/SDC1 and ANGPTL4/SDC4 communication components with RNA, latent states and model parameters fixed. **b** Mean velocity changes after removal of ANGPTL4, FN1 or CLDN1 communication components in the same target receiver cohorts; the point area indicates the fraction of receivers with positive incoming influence from the targeted signalling modules. **c** Mean changes in all six gene velocities after joint removal. **d** Full-model velocity (left) and changes after joint removal (right), with symmetric logarithmic colour scales shared across sections within each group. **e** Changes in the six-gene local spatial projection after joint removal of the targeted communication components. Unit arrows on the original tissue coordinates represent projected component-off minus full-model vectors, normalised after subtraction, and encode response direction only. **f** Paired full-to-component-off angles across 900 shared genes after scaling by gene-specific input standard deviations; points and horizontal lines indicate within-section medians and interquartile ranges. **g** Independent liver RNA responses to ANGPTL4 neutralisation or myeloid *Angptl4* deletion in LIDPAD mice at 8 and 12 weeks. Points represent mice, with five per group; bars indicate six-gene programme means. Exact two-sided permutation tests use Benjamini-Hochberg correction across four comparisons. **h** Independent gene-expression differences relative to LIDPAD controls, expressed as Δlog_2_(1 + CPM). Velocity changes in b-d denote component off minus full, in input standard deviation units with indicated multipliers. ND, normal diet; LP, LIDPAD; Ab, antibody; KO, myeloid deletion.

